# Modulating hydrogel architecture via cross-linker length for high-resolution tissue imaging and photochemical sectioning

**DOI:** 10.64898/2026.08.15.743111

**Authors:** Guang Yang, Wei Wang, Rahul Mitra, Ruixuan Gao

## Abstract

The recent development of Volumetric Imaging via Photochemical Sectioning (VIPS) has enabled nanoscale imaging of whole-mount tissue samples of virtually any size by embedding intact tissue in a photocleavable, superabsorbent hydrogel. However, the efficacy of sample embedding, imaging, and photochemical sectioning is fundamentally governed by the mechanical stiffness, structural stability, and photodegradation kinetics of the photocleavable hydrogel (PC-gel) polymer network. To elucidate the effect of the photosensitive crosslinker design on these critical properties, we synthesized a set of photocleavable crosslinkers (PCs) with varying polyethylene glycol (PEG) backbone lengths and prepared the corresponding PC-gels under a fixed monomer formulation and polymerization condition. We quantified and compared the viscoelastic properties of the formed PC-gels at their swollen states, and found that the crosslinker length markedly reshaped the PC-gel mechanics. In addition, we evaluated the light-triggered degradation of the PC-gels using both wide-field and spatially-controlled illumination. We found that PC-1000, PC-1500, and PC-2000 gels remained comparably photodegradable, all enabling on-demand, spatially confined decrosslinking under such illuminations. These results provide practical guidelines for modulating the crosslinker architecture of PC-gel polymer networks to achieve optimal physicochemical properties for whole-mount tissue imaging using VIPS.

## Introduction

Recent advances in hydrogel-based tissue clearing and expansion have transformed optical imaging of cellular and subcellular structures and spatial mapping of their molecular profiles across intact specimens.^1^ In hydrogel-based tissue embedding and expansion, biological specimens undergo in situ polymerization to form a sample-hydrogel composite, where biomolecules of interest are anchored to the hydrogel polymer networks via covalent or non-covalent conjugation.^2,3^ Crucially, the polymeric architecture of the embedding hydrogel dictates the physicochemical properties of the resulting tissue-hydrogel composite, including its structural homogeneity, mechanical resilience, and optical transparency.

When coupled with high-throughput volumetric imaging modalities, such as light-sheet fluorescence microscopy (LSFM), hydrogel-based tissue transformation provides a robust platform for molecular imaging of whole-mount tissue samples.^4^ The inherent advantages of this integration, which include enhanced effective resolution, high-speed volumetric scanning, and reduced photobleaching, enable the nanoscale reconstruction of intact tissue specimens with high speed and molecular fidelity.^5,6^ However, the increasing dimension of hydrogel-transformed samples often pushes the tissue volumes beyond the working distance of high-numerical-aperture objectives. Consequently, high-resolution imaging of large-scale biological specimens remains fundamentally constrained without physically sectioning the sample.

To overcome this limitation, we recently introduced Volumetric Imaging via Photochemical Sectioning (VIPS).^7^ For VIPS, we designed a photocleavable hydrogel (“PC-gel”) capable of rapid and spatially controlled photodegradation of embedded tissue volumes under single- or multi-photon excitation. This approach overcomes the abovementioned inherent imaging depth limitations by facilitating iterative imaging and exposure of deeper tissue layers through a non-contact sectioning process that minimizes sample damage.

To implement VIPS, PC-gel is synthesized by incorporating a photocleavable crosslinker (PC) in a polyacrylamide/sodium polyacrylate hydrogel network (**Figure 1A**). This original PC molecule features a ∼2 kDa polyethylene glycol (PEG) backbone end-functionalized with two photocleavable *o*-nitrobenzyl (ONB) ester moieties.^7^ Upon UV irradiation, the ONB ester groups undergo rapid photolysis, triggering crosslinker cleavage and consequent degradation of the hydrogel polymer networks. While the PEG backbone of PC provided sufficient hydrophilicity and biocompatibility, how its length and architecture dictate the mechanical and optical properties of the PC-gel for VIPS has not yet been systematically investigated.

**Figure 1.**
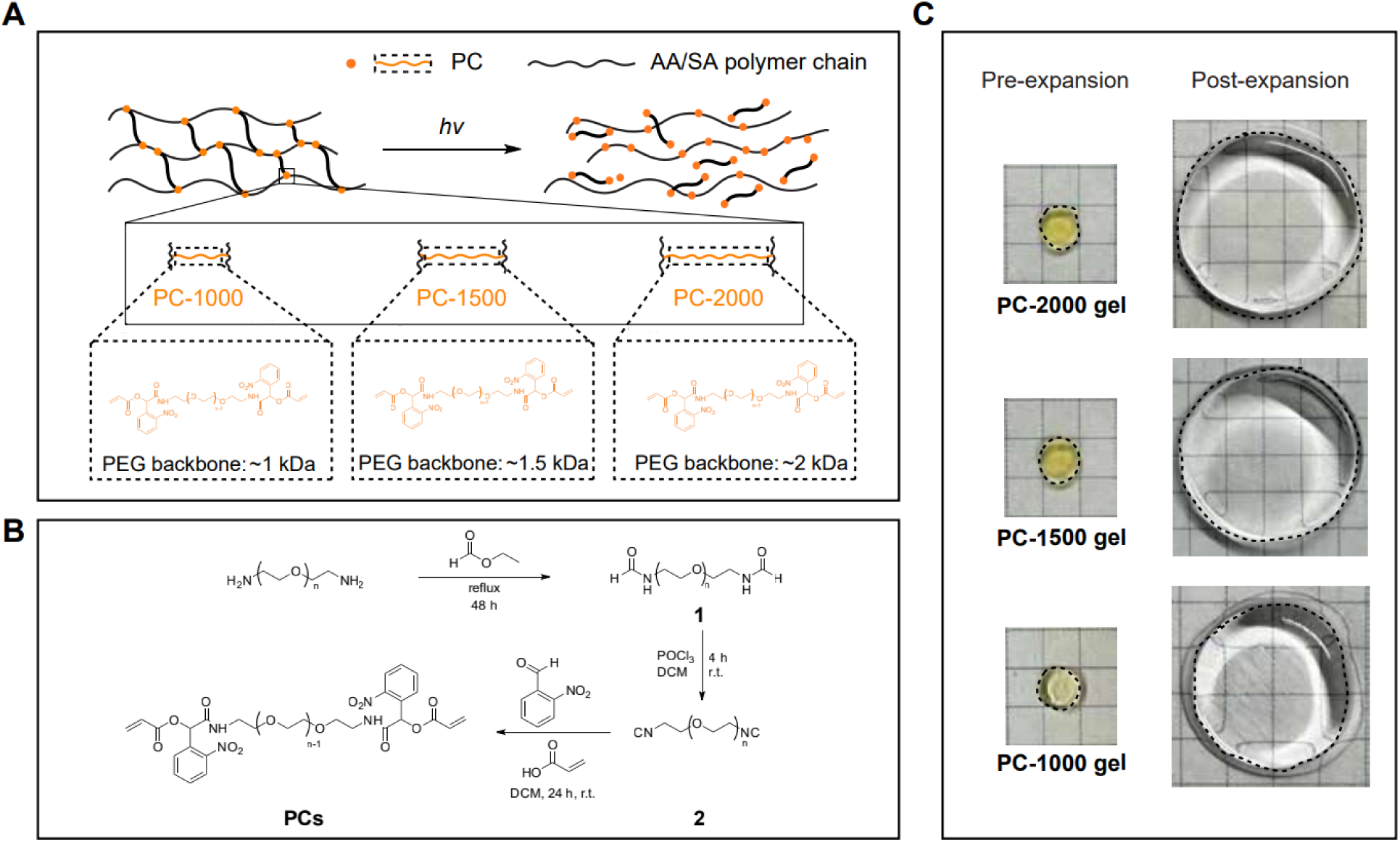
Design, synthesis, and expansion behavior of polyacrylamide/sodium polyacrylate hydrogels crosslinked with photocleavable crosslinkers (PC-gels) of varied lengths. (**A**) Schematic of acrylamide/sodium acrylate (AA/SA) polymer networks (black chains) crosslinked by PC (orange) bearing an *o*-nitrobenzyl (ONB) ester photolabile motif. UV irradiation photocleaves the PC linkage to allow controlled photodegradation of the photocleavable hydrogel (PC-gel) polymer networks. The enlarged dashed boxes show the representative PC-1000, PC-1500, and PC-2000 crosslinkers. (**B**) Synthetic route for PC derivatives (n = ∼2, ∼8, ∼20, ∼32, or ∼43). PEG diamine is converted to a bis-formamide intermediate (**1**), dehydrated to the corresponding diisocyanide (**2**) under POCl□ conditions, and subsequently subject to a Passerini reaction (with 2-nitrobenzaldehyde and acrylic acid) to add the ONB and acrylate termini, yielding polymerizable PCs. (**C**) Representative photographs of PC-gels (PC-1000, PC-1500, and PC-2000 gels) before (as-gelled, pre-expansion) and after (post-expansion) swelling. Grid size: 5 mm.

Previous studies have provided important insights into the potential influence of crosslinker architecture on the hydrogel polymer networks. For example, crosslinker length has been reported to affect gelation kinetics and mechanical properties of hydrogels.^8–13^ In a hyperbranched polyglycerol (hPG-NH ) hydrogel system crosslinked by PEG dialdehyde, reducing the PEG backbone length from 3 kDa to 1 kDa resulted in faster gelation and increased strength.^10^ Conversely, in thiol-ene hydrogel systems, longer PEG crosslinkers (3.5 kDa versus 1.5 kDa) were reported to promote more rapid gelation,^9^ highlighting that the relationship between crosslinker length and gelation behavior is system-dependent.

In hydrogel-based tissue transformation, smaller crosslinkers typically offer an advantage due to their enhanced diffusion into dense tissue specimens. However, reducing the PEG backbone length of PC may also compromise its aqueous solubility, as the hydrophobic influence of the two ONB ester groups becomes more pronounced. From a synthetic standpoint, the PEG chain length dictates the physical state and handling of the PC, as PEG transitions from a viscous liquid to a crystalline solid with increasing molecular weight. Consequently, establishing how the PEG backbone length modulates PC-gel polymerization, physicochemical attributes, and photochemical sectioning efficiency is of significant interest to researchers utilizing the VIPS approach (**Figure 1A**).

To investigate the effect of PEG backbone length on the PC-gel, we synthesized several PC derivatives with varying PEG backbone lengths: PC-200, PC-500, PC-1000, PC-1500, and PC- 2000. Each of these crosslinkers was then incorporated into a polyacrylamide/sodium polyacrylate hydrogel network to evaluate their ability to form a photodegradable and superabsorbent PC-gel. Of the five PC derivatives tested, three successfully formed stable PC-gels, which we further characterized by evaluating their expansion factors, rheological properties, and photodegradation behavior. Collectively, this work establishes a structure-property framework for selecting optimal photocleavable crosslinker length to achieve robust tissue gelation, imaging, and photochemical sectioning.

## Results and Discussion

### Synthesis of PC derivatives with varied PEG backbone lengths

For the initial implementation of VIPS, PC was synthesized with a PEG backbone length of ∼2 kDa (PC-2000) using polyethylene glycol bis(2-aminoethyl) ether with a number-averaged molecular weight (Mn) of ∼2000 g/mol (PEG-2000) as the precursor.^7^ We hypothesized that varying the PEG backbone length would alter the mechanical stiffness and the photodegradation kinetics of the PC-gel. Therefore, we obtained PEG diamine precursors with systematically shortened PEG backbone lengths, including PEG-200, PEG-500, PEG-1000, PEG-1500, and PEG-2000. These precursors were then subjected to a similar multi-step synthesis to yield PC derivatives, with stoichiometric ratios adjusted based on their molecular weights (**Figure 1B**). As a result, PCs with varying PEG backbone lengths were successfully synthesized, including PC- 200, PC-500, PC-1000, PC-1500, and PC-2000 (**Figure S5-S9**).

To isolate the specific effect of PEG backbone length on hydrogel properties, the molar concentration of PC was standardized across all experimental groups during PC-gel formation. In practice, determining the precise crosslinker molarity of PC derivatives was challenging because the purity of the as-synthesized products varied across the batches, and the PEG-derived impurities in the final products were difficult to remove completely.^14–16^ In addition, PEG-rich compounds were often hygroscopic,^17,18^ further increasing the uncertainty in the final yield and purity of the products.

To address this challenge, we used quantitative ¹H NMR (qNMR) to determine the batch-specific PC molarity prior to gelation. In this experiment, ^1^H NMR measurements of as-synthesized PC derivatives were performed in D_2_O to reflect the aqueous environment used for PC-gel formation. Acetone was selected as the internal standard because it is fully miscible with D_2_O, thus providing a single sharp ¹H resonance in a region that does not overlap the diagnostic PEG signals, and can be readily removed due to its volatility, minimizing the likelihood of carryover. The absolute amount of PC was then determined using the conserved benzylic methine (CH) at the acrylate–*o*-nitrobenzyl-PEG-amide junction, which is present in all PC derivatives and remains distinct from the PEG precursor resonances. Finally, each batch was measured in triplicate, and the average qNMR-derived molarity was used to prepare normalized crosslinker stock solutions to ensure identical PC molar concentrations in the final gelling solution.^19–21^ Quantitative acquisitions as well as the processing details and representative qNMR spectra are provided in the **Supporting Information** and **Figure S10-S14**.

Because aqueous solubility and microheterogeneity can, in principle, bias quantitative spectral integration in water-rich media, we further performed an orthogonal qNMR validation in an organic solvent system (CDCl ) using cyclohexane as the internal standard. Given the limited aqueous solubility and the incompatibility of PC-200 and PC-500 with PC-gel formation as described later, this cross-check was performed only for PC-1000, PC-1500, and PC-2000 (**Figures S15-S17**). As a result, the PC concentrations determined by CDCl /cyclohexane qNMR were consistent with those obtained in the D_2_O/acetone, confirming the reliability of the assigned PC stock-solution molarities.

### Synthesis of PC-gels with varied PC lengths

Prior to PC-gel synthesis, PC derivatives with different PEG backbone lengths (PC-200, PC-500, PC-1000, PC-1500, and PC-2000) were dissolved in water to prepare 1 M stock solutions. This concentration was matched with the standard crosslinker molarity used to prepare PC-gel in the VIPS study.^7^ Among the evaluated PC derivatives, PC-200 and PC-500 exhibited drastically poorer aqueous solubility, resulting in the formation of light-yellow precipitates. Both solutions were then filtered to remove the insoluble precipitates and polymerized in a 37 °C incubator. After gelation, no PC-gel was formed with the PC-200 stock solution, and only the central portion of the gelation chamber polymerized with the PC-500 stock solution. Upon swelling in water, the partially formed PC-500 gel exhibited an apparent ∼10-fold expansion factor but showed extremely weak mechanical integrity, rendering it unsuitable for VIPS or other imaging applications. We note that this apparent high expansion factor of PC-500 gel should be interpreted differently from the trend observed for fully formed PC-gels. Because a large fraction of PC-500 precipitated during gelation-solution preparation, the actual amount of PC-500 available for polymerization was likely much lower than the nominal amount added. As a result, the gel formation proceeded with far fewer crosslinkers, yielding a weaker gel with an unusually high swelling rate.

Unlike the shorter-chain PCs (PC-200 and PC-500), the longer-chain PCs (PC-1000, PC-1500, and PC-2000) could be prepared as aqueous stock solutions and used for PC-gel formation. However, their behavior in the gelation mixture showed distinct variations across the three types of PCs. PC-1000, and to a lesser extent PC-1500, became cloudy or partially precipitated within the stock solution, whereas PC-2000 remained soluble. The PC stock solutions were subsequently used to prepare PC-gels with qNMR-normalized crosslinker molarities (**Supporting Information**). After swelling in water, PC-1000, PC-1500, and PC-2000 gels showed linear expansion factors of ∼3.3 ± 0.3-fold, ∼3.8 ± 0.2-fold, and ∼4.5 ± 0.3-fold, respectively (mean ± SD, n = 3 independently prepared gels; **Figure 1C** and **Table S1**). This trend may be attributed to the increasing hydrophilicity and flexibility of PC as its PEG backbone length increases. Both factors can facilitate water incorporation, allowing the polymer network to swell more extensively.

### Mechanical properties of PC-gels with varied PC lengths

The mechanical properties of hydrogels are important for hydrogel-based sample transformation because mechanically robust gels are easier to handle and more resistant to deformation during sample processing, handling, and imaging. To characterize the mechanical properties of PC-gels with different crosslinker lengths, we performed oscillatory rheological measurements on fully swollen gels. Because these water-swollen gels were sensitive to compression and strain-dependent softening, the measurement conditions were selected based on preliminary trials to obtain stable and reproducible responses (**Supporting Information**). During measurement, the PC-gels were maintained in a defined water reservoir to prevent dehydration and minimize modulus changes caused by water loss or displacement. Frequency sweeps were performed at a fixed low shear strain of 0.01% from 0.1 to 10 rad/s, followed by amplitude sweeps from 0.01% to 50% shear strain at a fixed angular frequency of 10 rad/s. Three independently prepared gels were measured for each formulation, with the replicate datasets provided in the **Supporting Information** (**Figures S2-S4**).

Representative frequency-sweep data are shown in **Figure 2A–C**. Across the tested angular-frequency range, the storage modulus was generally higher than the loss modulus for all three swollen PC-gel formulations. The overall stiffness followed the order PC-2000 > PC-1500 > PC- 1000, with PC-2000 showing the highest storage modulus and PC-1000 the lowest. Representative amplitude-sweep data are shown in **Figure 2D–F**. All three swollen PC-gels showed strain-dependent softening as the applied shear strain increased. PC-2000 showed the highest low-strain modulus, followed by PC-1500 and PC-1000. These results indicate that PC- 2000 forms the stiffest swollen PC-gel under the conditions tested, whereas PC-1000 remains the softest formulation across the three gels tested.

**Figure 2.**
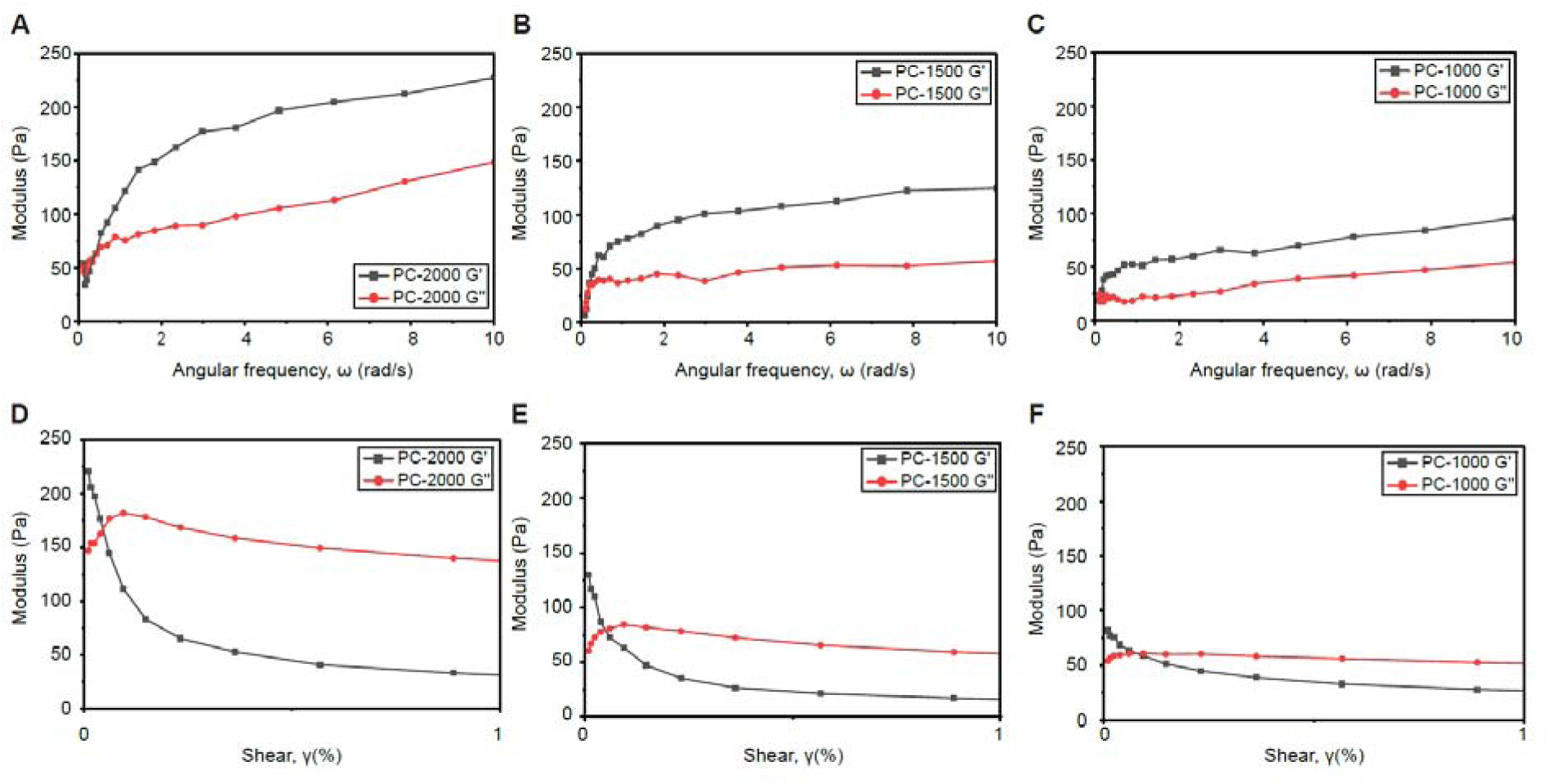
Oscillatory rheological characterization of swollen PC-gels with varied crosslinker lengths. (**A–C**) Representative frequency-sweep measurements of swollen PC-2000, PC-1500, and PC-1000 gels, respectively. Frequency sweeps were performed from 0.1 to 10 rad/s at a fixed shear strain of 0.01%. (**D–F**) Representative amplitude-sweep measurements of the corresponding swollen PC-gels. Amplitude sweeps were performed from 0.01 to 50% shear strain at a fixed angular frequency of 10 rad/s; the low-strain region from 0 to 1% shear strain is shown for clarity. Storage modulus (G’) and loss modulus (G”) are shown. The representative curves shown here correspond to one independently prepared gel from each formulation. Replicate measurements from three independently prepared gels are provided in the **Supporting Information**.

The lower modulus of PC-1000, and to a lesser extent PC-1500, may be related to their limited aqueous solubility during gel preparation. The cloudiness and occasional precipitation observed with these formulations suggest that these PCs may undergo local aggregation or less uniform incorporation into the polymer network, leading to softer, more heterogeneous gels. The smaller expansion factors observed for PC-1000 and PC-1500 likely also reflect less uniform gelation and reduced crosslinker incorporation efficiency.

### Photodegradation of PC-gels with varied crosslinker lengths

To successfully perform whole-mount tissue imaging and reconstruction using VIPS, a rapid, complete, and spatially precise photochemical degradation of PC-gel is required. Although no previous studies have reported the direct effects of PEG backbone lengths on photolysis kinetics, longer PEG chains may introduce increased physical entanglements within the polymer network, thereby potentially slowing the photodegradation by restricting network disassembly. To test this, we photodegraded bulk PC-gels using PC-1000, PC-1500, and PC-2000 as the crosslinker. Following expansion in water, the gels were exposed to ultraviolet (UV) illumination at 365 nm to induce photochemical degradation as previously described.^7^ As a result, all three PC-gels were fully photodegraded within 3.5 hours. The photodegradation time for the PC-2000, PC-1500, and PC-1000 gels was ∼3, ∼3.5, and ∼3 hours, respectively (**Figure 3A**), indicating that PEG chain lengths only have a minor impact on the bulk gel photodegradation kinetics. The non-monotonic trend, where the photodegradation time first increases and then decreases with increasing PEG length, suggests that additional factors may have affected the bulk photodegradation measurements. One possible explanation is that the PC-1000 gel would have intrinsically required a comparatively longer time for complete bulk-gel photodegradation (e.g., 3.5 hours or more). However, its lower stiffness may have facilitated apparent gel disintegration, resulting in an overall bulk-gel photodegradation time similar to that of the PC-2000 gel.

**Figure 3.**
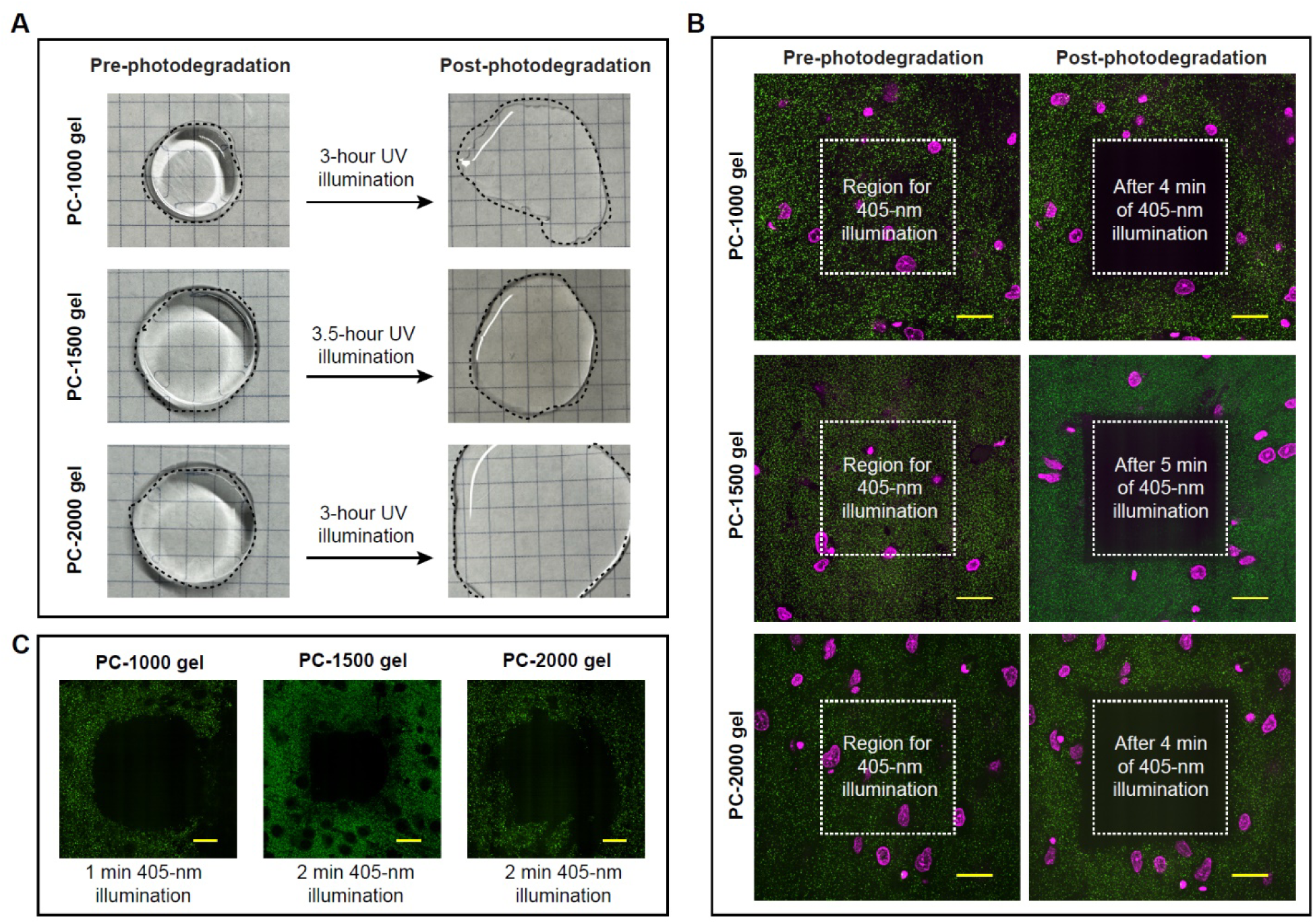
Photodegradation of PC-gels with varied crosslinker lengths. (**A**) Bulk photodegradation of PC-1000, PC-1500 and PC-2000 gels under a UV lamp. Photographs of PC-gels pre- (left) and post-illumination (right) are shown, with the gel contour indicated by a dashed line. Grid size: 5 mm. (**B**) Photodegradation of PC-1000-, PC-1500- and PC-2000-gel-embedded tissue using spatially confined confocal illumination. Pre- (left) and post- (right) 405-nm laser illumination images of mouse brain tissue fluorescently labeled for postsynaptic protein (Homer 1, Alexa Fluor 488, green) and nuclei (SiR-DNA, magenta) and expanded with the PC-gels are shown. Scale bars, 100 μm (post-expansion scale). (**C**) Early onset photodegradation of PC-1000-, PC-1500- and PC-2000-gel-embedded tissue using the spatially confined confocal illumination. Similarly, fluorescently labeled mouse tissue as **B** was used. Scale bars, 100 μm (post-expansion scale).

To further assess the efficacy of photodegradation, mouse brain tissue labeled with anti-Homer 1 antibodies and SiR-DNA was embedded within each PC-gel type. Here, SiR-DNA was used instead of DAPI for the nuclear counterstaining to avoid unwanted photodegradation during DAPI imaging. Following gelation and expansion in water, selective-area photodegradation experiments were performed using a confocal microscope equipped with a 405 nm laser (**Figure 3B**).^22–24^ We note that the observed loss of the sample fluorescence within the illuminated area can be attributed to photodegradation and not photobleaching, as validated in the previous VIPS study.^7^ Across the three PC-gels, no significant differences in the photodegradation spatial accuracy were observed. In fact, all formulations exhibited highly localized and well-defined sample photodegradation after several minutes of laser illumination.

The photodegradation onset showed slight differences according to the PC length. While the PC- 1000 gels began photodegrading after ∼1 minute of the 405 nm laser exposure, the PC-1500 and PC-2000 gels required ∼2 minutes (**Figure 3C**). The total photodegradation times also followed a similar trend as the bulk photodegradation experiments: ∼4 minutes for the PC-1000 and PC- 2000 gels, and ∼5 minutes for the PC-1500 gel. Combined, these results suggest that (i) the PEG backbone length does not substantially alter the intrinsic photocleavage kinetics of the formed PC-gels, (ii) the observed subtle increase in photodegradation onset and completion time from the PC-2000 gel to the PC-1500 gel may be caused by the lower expansion factor of the PC-1500 gel, and (iii) the non-monotonic trend mentioned above between the PC length and photodegradation time may be attributed to the PC-length-dependent stiffness, which dictates the threshold at which macroscopic softening becomes detectable (i.e., the PC-1000 gel showing lower gel stiffness relative to the PC-1500 and PC-2000 gels).

## Conclusion

In this study, we synthesized a set of PCs with varied PEG backbone lengths (PC-200, PC-500, PC-1000, PC-1500, and PC-2000) and systematically evaluated how the crosslinker length impacts PC-gel formation and its properties for sample handling, imaging, and photochemical sectioning. PC-200 and PC-500 exhibited limited aqueous solubility under targeted crosslinker concentrations, leading to failed (PC-200) or incomplete (PC-500) PC-gel formation.

To ensure a fair comparison across the remaining PC derivatives, the PC concentration was normalized by qNMR prior to gelation, and an orthogonal solvent system was used to validate the molarity assignments. As a result, all three longer PCs (PC-1000, PC-1500, and PC-2000) successfully formed stable PC-gels swellable in water, with their linear expansion factors being ∼3.3 ± 0.3-fold, ∼3.8 ± 0.2-fold, and ∼4.5 ± 0.3-fold, respectively. At the same molar crosslinker concentration, longer PCs may have increased the hydrophilic PEG contents and introduced longer and flexible crosslinkers within the PC-gel polymer network, thereby promoting water uptake and swelling.

Oscillatory rheological measurements of the fully swollen PC-gels showed that the PC-2000 exhibited the highest storage modulus, followed by the PC-1500 and PC-1000 gels. The PC-1000 remained the softest formulation. This lower modulus may be attributed to its limited aqueous solubility during gel preparation, which could have led to less and nonuniform crosslinker incorporation and a softer, more heterogeneous polymer network.

Finally, all three PC-gel formulations (PC-1000 gel, PC-1500 gel, and PC-2000 gel) exhibited satisfactory photochemical degradation properties, both in bulk and within embedded biological tissues. Under wide-field UV illumination, complete photodegradation was observed by ∼3.0 hours for the PC-1000 and PC-2000 gels, and by ∼3.5 hours for the PC-1500 gel. Under spatially confined 405 nm confocal illumination, all PC-gels exhibited similarly high spatial confinement in their photodegradation, with only minimal differences observed for the onset and overall photodegradation times. Overall, these results revealed no significant differences in the photodegradation kinetics and spatial control across the three PC-gels.

For hydrogel-based tissue embedding and expansion, efficient penetration of the gelling solution into dense biological tissues is critical to ensure uniform gelation throughout the specimen. In principle, larger crosslinkers diffuse more slowly across thick tissues due to their increased molecular size. However, this limitation can potentially be overcome by extending the incubation time of the gelling solution with biological samples before polymerization. For example, in the previous VIPS study,^7^ extending the pre-polymerization incubation time of the gelling solution from hours to days enabled successful gelation of whole mouse olfactory bulbs that were several millimeters thick. Furthermore, if even larger tissue specimens need to be gelled (e.g., the whole mouse brain), switching to an alternative initiator system can allow for an even longer pre-polymerization incubation period, providing additional time for the gelling solution to fully penetrate the tissue before polymerization begins.

Taken together, our results show that PC-2000 represents a balanced choice for VIPS because it exhibits (i) adequate aqueous solubility and robust PC-gel formation, (ii) high post-expansion mechanical integrity (e.g., the highest G’ among all the PC-gels tested), (iii) satisfactory photodegradation performance. On the other hand, PC-1000 suffers from the issue of lower aqueous solubility, which can result in a compromised mechanical property. PC-1500 remains a viable option for VIPS when shorter gelling solution incubation time is needed, while the PC- 1000 gel may exhibit a slightly lower mechanical integrity due to the lower aqueous solubility as compared to the PC-2000 gel. Collectively, our results establish a structure-property relationship between PC-gel architecture and its crosslinker length and serve a practical guide to selecting the PC design to balance gelation robustness, expansion behavior, mechanical properties, and photochemical sectioning efficacy for VIPS.

## Author Contributions

R.G., W.W., and G.Y. conceived the project and supervised the work; G.Y. performed the synthesis of compounds and participated in all experimental processes; W.W. participated in all experimental processes except for the synthesis; R.M. contributed to the gelation experiments, image acquisition, and characterization studies; G.Y., W.W., and R.M. contributed to the characterization and data analysis; G.Y., W.W., and R.M. prepared the figures. W.W. and G.Y. wrote the original draft. R.G. validated, reviewed, and edited the writing. All authors have reviewed and approved the final version of the manuscript. G.Y. and W.W. contributed equally to this work.

## Competing Interests

The authors declare the following competing financial interest(s): R.G. is a co-inventor of multiple patents related to expansion microscopy and VIPS. The other authors declare no competing financial interests.

## Acknowledgments

We thank the Analytical BioNanoTechnology Equipment Core (ANTEC), Center for Regenerative Nanomedicine at Northwestern University, and the Chicago Biomedical Consortium (CBC) for access to rheometry instrumentation. We also thank Hiroaki Sai (Acting Director of ANTEC) for assistance and training on the rheometer. We thank Daniel McElheny (Director of NMR Facilities, University of Illinois Chicago) for assistance with qNMR measurements and analysis. R.G. acknowledges the National Institutes of Health (grant number: UG3MH126864), University of Illinois Chicago Startup Fund, the Searle Scholars Program, and the McKnight Technological Innovations in Neuroscience Award for funding support.

