## Supplemental Information for "Modulating hydrogel architecture via cross-linker length for high-resolution tissue imaging and photochemical sectioning"

**Table of contents**

1. General synthetic methods
2. Synthetic procedure and characterization
3. Biological sample preparation
4. Basic gelation protocol
5. Sample imaging and photodegradation
6. Rheometry
7. Quantitative NMR analysis
8. NMR spectra
9. References

### 1. General synthetic methods

Unless stated otherwise, all commercially obtained reagents and solvents were used as received without further purification. All solvents used for spectroscopy were of spectroscopy grade. Reactions were conducted in oven-dried glassware. Unless stated otherwise, $^{1}H$-NMR spectra were obtained using a 500 MHz NMR spectrometer (Bruker). Data from $^{1}H$-NMR spectroscopy are reported as chemical shifts (δ ppm) with the corresponding integration values. Coupling constants (*J*) are reported in hertz (Hz). Standard abbreviations indicating multiplicity are as follows: *s* (singlet), *b* (broad), *d* (doublet), *t* (triplet), *q* (quartet), *m* (multiplet), and *virt* (virtual). Unless stated otherwise, all chemicals and reagents were obtained from Millipore Sigma.

#### 1.1 General synthetic route

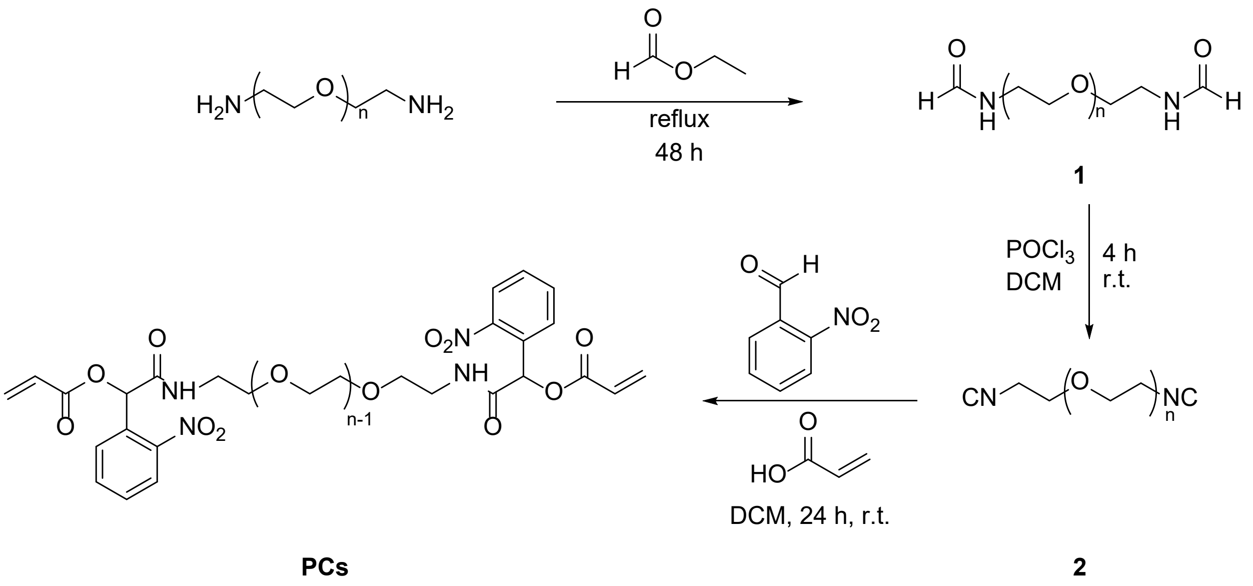


**Scheme S1.** General synthetic route for photocleavable cross-linker (**PC**) derivatives. n = ~2, ~8, ~20, ~32, or ~43.

### 2. Synthetic procedure and characterization

No unexpected or unusually high safety hazards were encountered. Phosphoryl chloride ($POCl_3$) and volatile organic solvents were handled in a chemical fume hood using appropriate personal protective equipment.

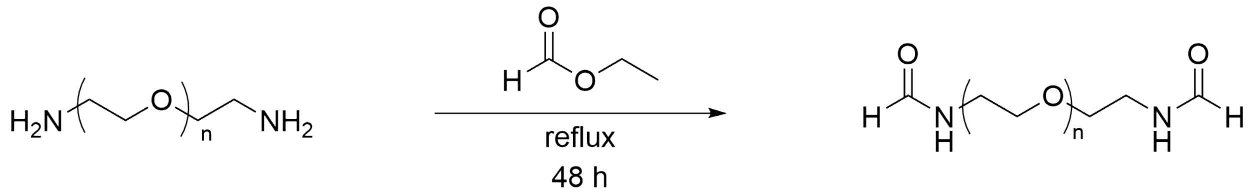


1

**Scheme S2.** General synthetic route for polyethylene glycol (PEG)-based bis(2-formamidoethyl) ether homologs (**1**). The structure shown represents the general form of the products, where n = ~2, ~8, ~20, ~32, or ~43 corresponding to the PEG-200, PEG-500, PEG-1000, PEG-1500, and PEG-2000 backbones, respectively.

#### 2.1 Synthesis of polyethylene glycol bis(2-formamidoethyl) ether homologs (1)

Polyethylene glycol bis(2-aminoethyl) ethers (with PEG-200, PEG-500, PEG-1000, PEG-1500, and PEG-2000 backbones; Biopharma PEG Scientific Inc.) were converted to the corresponding polyethylene glycol bis(2-formamidoethyl) ether homologs (**1**) following a modified literature procedure (**Scheme S2**).[1–3]

**For liquid-state PEGs (PEG-200 and PEG-500):**

The liquid diamine precursor (200 mg) was first mixed with a small amount of ethyl formate (2-3 mL) to form a homogeneous solution, which was then transferred to a reaction vial under a nitrogen atmosphere. The original vial was rinsed with additional ethyl formate to bring the total solvent volume to 8 mL. The mixture was ultrasonicated briefly (5 to 10 min) to ensure complete dissolution, then heated to reflux (≈ 55 °C) with magnetic stirring for 48 h. After completion, the reaction mixture was cooled to room temperature, and the solvent was removed under reduced pressure using a rotary evaporator (N-1300, EYELA; water bath set to 32 °C). The polyethylene glycol bis(2-formamidoethyl) ether homologs **(1)** were obtained as clear, transparent, colorless, viscous liquids and used directly in subsequent reactions.

**For solid-state PEGs (PEG-1000, PEG-1500, and PEG-2000):**

The diamine precursor (200 mg) was added to ethyl formate (8 mL) under a nitrogen atmosphere, and the mixture was gently heated with intermittent ultrasonication until a clear, homogeneous solution was obtained. The solution was then maintained at reflux (≈ 55 °C) with vigorous stirring for 48 h. After the reaction was complete, the mixture was cooled to room temperature, and the solvent was removed under reduced pressure using a rotary evaporator (N-1300, EYELA; water bath set to 42 °C) to afford the corresponding polyethylene glycol bis(2-formamidoethyl) ether homologs (**1**) as white to off-white waxy solids. Reaction progress was monitored by thin-layer chromatography (TLC) using a dichloromethane (DCM)/hexane (4:6, v/v) solvent system, with the disappearance of the amine precursor indicating completion of the reaction. All crude materials were used directly in subsequent reactions without further purification.

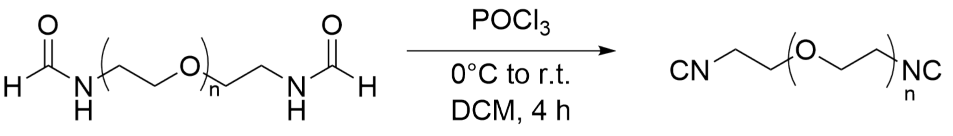


**1** **2**

**Scheme S3.** Synthesis of polyethylene glycol bis(2-isocyanoethyl) ether homologs (**2**) from polyethylene glycol bis(2-formamidoethyl) ether homologs (**1**) using $POCl_3$-mediated dehydration. n = ~2, ~8, ~20, ~32, or ~43.

**2.2 Synthesis of polyethylene glycol bis(2-isocyanoethyl) ether homologs (2)**

Polyethylene glycol bis(2-formamidoethyl) ether homologs (**1**; with **PEG-200**, **PEG-500**, **PEG-1000**, **PEG-1500**, or **PEG-2000** backbones) were dehydrated with $POCl_3$ under basic conditions to afford the corresponding polyethylene glycol bis(2-isocyanoethyl) ether homologs (**2**) (**Scheme S3**).

Briefly, in an oven-dried, septum-sealed flask, the polyethylene glycol bis(2-formamidoethyl) ether homologs intermediate (**1**) (1.0 equiv) was dissolved in anhydrous DCM (15 mL) under a nitrogen atmosphere, and the solution was cooled in an ice bath for 10 min. Triethylamine (TEA, 4.0 equiv) was added, and the mixture was stirred vigorously. In a separate oven-dried vessel, $POCl_3$ (1.5 equiv) was dissolved in anhydrous DCM (3 mL), and this solution was added dropwise over 20 min to the cold reaction mixture. Gas evolution and gradual color change were typically observed. The addition rate was adjusted to maintain the internal temperature around 0 °C and to prevent pressure buildup. After the addition was complete, the mixture was maintained at 0 °C for 2 min, then allowed to warm to room temperature, and stirred for 4 h. Upon completion, the reaction mixture was cooled in an ice bath and quenched with chilled saturated $K_2CO_3$ solution. After gas evolution subsided, the mixture was transferred to a separatory funnel and allowed to separate into layers (vigorous shaking was avoided). The organic phase was collected, and the aqueous phase was extracted with DCM three times. The combined organic extracts were washed with saturated NaCl (brine) once or more (repeated as needed), dried over anhydrous $Na_2SO_4$, filtered, and concentrated under reduced pressure to a concentrated solution (complete drying was avoided). Product **2** was precipitated by adding the concentrated solution dropwise into cold diethyl ether. The precipitate was collected by centrifugation, the supernatant was decanted, and the solid was dried under vacuum to afford the polyethylene glycol bis(2-isocyanoethyl) ether homologs (**2**) as a viscous liquid for **PEG-200** and **PEG-500**, or a red-orange to yellow solid for the rest of the products. Owing to the moisture sensitivity of isocyanides, the crude product was used immediately in the subsequent step without further purification.

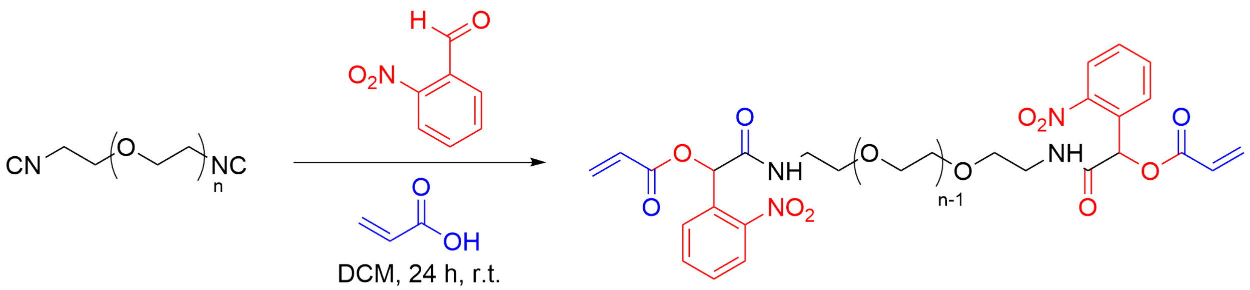


**2** **PCs**

**Scheme S4.** Synthesis of PEG-based photocleavable cross-linker (**PC**) derivatives from polyethylene glycol bis(2-isocyanoethyl) ether homologs (**2**). n = ~2, ~8, ~20, ~32, or ~43.

### 2.3 Synthesis of PEG-based photocleavable cross-linker (PC) derivatives

The polyethylene glycol bis(2-isocyanoethyl) ether homologs (**2**; with **PEG-200**, **PEG-500**, **PEG-1000**, **PEG-1500**, and **PEG-2000** backbones; 1.0 equiv) were converted to the corresponding PEG-based photocleavable cross-linker (PC) derivatives (**PC-200**, **PC-500**, **PC-1000**, **PC-1500**, and **PC-2000**) through a three-component Passerini reaction with 2-nitrobenzaldehyde and acrylic acid (**Scheme S4**).

Briefly, the polyethylene glycol bis(2-isocyanoethyl) ether intermediates (**2**; 1.0 equiv), 2-nitrobenzaldehyde (3.0 equiv), and acrylic acid (3.5 equiv) were dissolved in anhydrous dichloromethane (DCM) under a nitrogen atmosphere. The reaction mixture was stirred at room temperature (≈ 25 °C) in the dark for 24 h. Extending the reaction beyond 24 h resulted in the formation of colored impurities and was therefore avoided. Upon completion, DCM was removed under reduced pressure, yielding a yellow, oily residue, which was dissolved in a minimal amount of ethyl acetate. Petroleum ether or hexane was gradually added until precipitation occurred, and the mixture was then cooled in an ice bath for 30 min. The resulting precipitates were collected by filtration, washed with cold petroleum ether or hexane, and dried under vacuum overnight, affording the as-synthesized PC derivatives (**PC-200, PC-500, PC-1000, PC-1500, and PC-2000**) as yellow solids. For gelation, the as-synthesized PCs were dissolved in double-distilled water to a concentration of 1 M. The aqueous PC stock solutions were then passed through a 0.22 μm PTFE syringe filter to remove insoluble precipitates, if any.

When further purification was needed, the as-synthesized PC derivatives were dissolved in deionized water to a concentration of ~25 mg/mL, and purified by dialysis using cellulose-based membranes/devices with nominal molecular weight cut-offs (MWCOs) selected to match the target product size. **PC-1500** and **PC-2000** were dialyzed against deionized water using Spectra/Por 6 tubing (MWCO: 1 kDa; 132638T, Repligen), while **PC-1000** was purified using a Pur-A-Lyzer Midi dialysis kit (MWCO: 1 kDa; PURD10005-1KT, Millipore Sigma). **PC-500** and **PC-200** were purified using low-MWCO Float-A-Lyzer G2 CE dialysis devices (MWCO: 0.1-0.5 kDa; 131054T, Repligen). In each case, the MWCO was selected to facilitate removal of low-molecular-weight impurities while minimizing loss of the desired PC product. Dialysis was performed against deionized water with multiple water exchanges over 12-24 h. The aqueous solution was then passed through a 0.22 μm PTFE syringe filter to remove insoluble precipitates and concentrated under reduced pressure. The product was reprecipitated from ethyl acetate using petroleum ether or hexane, affording the purified PC derivatives (**PC-200**, **PC-500**, **PC-1000**, **PC-1500**, and **PC-2000**) as yellow-white powders. The purified PC derivatives were only used for selected experiments.

**PC-200.** *$^1H$ NMR* (500 MHz, $CDCl_3$) δ 7.98 (*d*, *J* = 8.8 Hz, 2H, Ar–H), 7.75 (*d*, *J* = 8.0 Hz, 2H, Ar–H), 7.55 (*m*, 2H, Ar–H), 7.09 (*s*, 1H, Ar–H), 6.71 (*s*, 1H, $CH=CH_2$), 6.48 (*d*, *J* = 16.8 Hz, 1H, $CH=CH_2$), 6.20 (*dd*, *J* = 16.8, 10.4 Hz, 1H, $CH=CH_2$), 5.94 (*d*, *J* = 10.4 Hz, 1H, $CH=CH_2$), 3.60 (br *s*, PEG–$CH_2$).

**PC-500.** *$^1H$ NMR* (500 MHz, $CDCl_3$) δ 7.98 (d, *J* = 8.8 Hz, 2H, Ar–H), 7.73 (d, *J* = 8.0 Hz, 2H, Ar–H), 7.64–7.50 (m, 4H, Ar–H),7.04 (s, 1H, Ar–H), 6.73 (s, 1H, $CH=CH_2$), 6.50 (d, *J* = 16.8 Hz, 1H, $CH=CH_2$), 6.24–6.18 (m, 2H, $CH=CH_2$), 5.95 (d, *J* = 10.4 Hz, 1H, $CH=CH_2$), 3.61 (*br s*, PEG–$CH_2$).

**PC-1000.** *$^1H$ NMR* (500 MHz, $CDCl_3$) δ 7.97 (*d*, *J* = 8.8 Hz, 2H, Ar–H), 7.72 (*d*, *J* = 8.0 Hz, 2H, Ar–H), 7.62 (*s*, 1H, Ar–H), 7.50 (*d*, *J* = 8.0 Hz, 1H, Ar–H), 7.02 (*s*, 1H, Ar–H), 6.72 (*d*, *J* = 16.8 Hz, 1H, $CH{=}CH_2$), 6.49 (*d*, *J* = 16.8 Hz, 1H, $CH{=}CH_2$), 6.20 (*dd*, *J* = 16.8, 10.4 Hz, 1H, $CH{=}CH_2$), 5.93 (*d*, *J* = 10.4 Hz, 1H, $CH{=}CH_2$), 3.60 (*br s*, PEG–$CH_2$).

**PC-1500.** *$^1H$ NMR* (500 MHz, $CDCl_3$) δ 7.99 (*d*, *J* = 8.8 Hz, 2H, Ar–H), 7.74 (*d*, *J* = 8.0 Hz, 2H, Ar–H), 7.65–7.48 (*m*, 5H, Ar–H), 6.95 (*s*, 1H, Ar–H), 6.73 (*s*, 1H, $CH{=}CH_2$), 6.51 (*d*, *J* = 16.8 Hz, 1H, $CH{=}CH_2$), 6.24–6.20 (*m*, 2H, $CH{=}CH_2$), 5.95 (*d*, *J* = 10.4 Hz, 1H, $CH{=}CH_2$), 3.61 (*br s*, PEG–$CH_2$).

The NMR data for **PC-2000** was consistent with those previously reported.[2]

### 3. Biological sample preparation

All mouse procedures were conducted in compliance with the U.S. National Institutes of Health Guide for the Care and Use of Laboratory Animals and were approved by the University of Illinois Chicago Animal Care Committee.

#### 3.1 Mouse brain slices

9-week-old C57BL/6 mice were anesthetized with a ketamine/xylazine cocktail (ketamine hydrochloride injectable solution, 100 mg/mL, Covetrus; AnaSed/xylazine sterile solution, 20 mg/mL, Akorn) and were transcardially perfused with 4% (w/v) paraformaldehyde (PFA) in 1× PBS (10 mL). Brains were carefully dissected from the skull, post-fixed in 4% (w/v) PFA in 1× PBS at 4 °C for 1 day, and then stored in sucrose buffer (20% sucrose in 1× PBS) at 4 °C. Fixed brains were sectioned with a vibratome (VT1200S, Leica) to obtain ~50 μm coronal slices. All sections were stored in 1× PBS containing 0.03% (w/v) sodium azide at 4 °C until use.

For immunostaining, ~50 μm brain slices were permeabilized with 0.1% (w/v) Triton X-100 in 1× PBS for 15 min and were incubated in blocking buffer [5% (v/v) normal goat serum (NGS) and 0.1% (w/v) Triton X-100 in 1× PBS] at room temperature for >6 h. Slices were then incubated with a primary antibody solution (1:200 in blocking buffer) at 4 °C for 2 days [(rabbit anti-Homer1, 160003, Synaptic Systems), (rabbit anti-Neurofilament 200, N4142-.2ML, Millipore Sigma)]. After primary incubation, slices were washed with blocking buffer four times (30 min per wash) and were incubated with an Alexa Fluor 488-conjugated secondary antibody (1:200 in blocking buffer; goat anti-rabbit Alexa Fluor 488, A11008, Thermo Fisher) at 4 °C for 1 day. Finally, slices were washed with 1× PBS four times (30 min per wash), and stored in 1× PBS at 4 °C prior to the subsequent gelation procedure.

### 4. Basic gelation protocol

Double-distilled water was obtained from a Milli-Q IQ 7000 system.

#### 4.1 Photocleavable hydrogel ("PC-gel")

PC-gel-embedded tissue specimens were generated following a reported procedure with minor adjustments[2]. Briefly, fixed and immunolabeled sections were incubated with acryloyl-X, SE (AcX) (0.1 mg/mL in 1× PBS) at room temperature overnight. Excess AcX was removed by rinsing the tissues in 1× PBS three times (15 min per rinse). For monomer loading, samples were transferred into a prechilled monomer formulation composed of 1× PBS, 2 M NaCl, 8.625% (w/v) sodium acrylate, 2.5% (w/v) acrylamide, and 4.980% (w/v) as-synthesized PC, and were maintained at 4 °C overnight to ensure thorough infiltration. Immediately prior to casting, polymerization reagents were introduced by adding concentrated stock solutions of APS (10% w/v), TEMED (10% w/v), and 4-hydroxy-2,2,6,6-tetramethylpiperidin-1-oxyl (4HT; 0.5% w/v) to final concentrations of 0.2% (w/v), 0.2% (w/v), and 0.01% (w/v), respectively, to afford the gelling mixture. The samples were then placed into freshly prepared gelling solution and were incubated at 0 °C for 30 min to limit premature gelation while promoting uniform initiator distribution. The gelling solutions and tissue samples were not actively deoxygenated by nitrogen purging or vacuum degassing. The samples were subsequently arranged in gelation chambers matched to the desired geometry, and gelation was carried out for 2 h in a humidified 37 °C incubator to yield photocleavable hydrogels (PC-gels). After gelation, the tissue-gel composites were digested at room temperature for 12 h in digestion buffer containing Proteinase K (final concentration of Proteinase K: 8 U/mL). The samples were washed with 1× PBS at room temperature and stained with SiR-DNA (CY-SC007, Cytoskeleton) according to the manufacturer's instructions. Finally, the samples were washed several times with pure water for expansion and were ready for subsequent imaging and photochemical sectioning experiments.

Purified PCs were used only for the expansion isotropy analysis experiments, which followed the same gelation protocol above. The gelled samples were not stained with SiR-DNA, and the pre-expansion images were acquired before the Proteinase K digestion step.

#### 4.2 Expansion factor measurement

Blank PC-gels were prepared using the formulation and polymerization conditions described in Section **4.1**, except that no tissue was embedded and no digestion step was performed. After polymerization, the gels were transferred to excess double-distilled water and allowed to swell until their dimensions no longer changed. The linear expansion factor was calculated as the post-expansion gel diameter divided by the corresponding pre-expansion diameter. Three independently prepared gels were measured for each formulation. Results are reported as mean ± standard deviation (SD).

**Table S1. Linear expansion factors of PC-gel with varied cross-linker lengths.** Values are provided as the linear expansion factor (fold), with n = 3 independently prepared gels per formulation.

| PC used for gelation | Replicate 1 | Replicate 2 | Replicate 3 | Mean ± SD |
|---|---|---|---|---|
| PC-1000 | 3.0 | 3.4 | 3.5 | 3.3 ± 0.3 |
| PC-1500 | 3.6 | 3.8 | 4.0 | 3.8 ± 0.2 |
| PC-2000 | 4.4 | 4.8 | 4.3 | 4.5 ± 0.3 |

### 5. Sample imaging and photodegradation

#### 5.1 Bulk photodegradation

Blank PC-gels were prepared following the protocol described in Section **4.1**, except that no tissue was embedded and no digestion step was performed. A round gelation chamber was used for gel formation. The gels were then expanded in pure water and subsequently transferred to a UV chamber (365 nm, 60 W; PC-60-DJ, Phrozen Tech Co., Ltd.) for bulk photodegradation. Gel integrity was assessed at 0.5 h intervals during illumination, and the complete photodegradation time was recorded with this sampling resolution.

#### 5.2 Confocal microscopy imaging and photodegradation

A spinning-disk confocal system (CSU-W1, Yokogawa Electric Corp.) integrated with an Eclipse Ti2-E microscope body (Nikon) was used for both imaging and photodegradation of tissue samples. Specifically, imaging data were acquired using a 488 nm and 640 nm solid-state laser, a sCMOS camera (ORCA-Fusion BT C15440, Hamamatsu), and a CFI Apo Lambda S LWD 20× (NA 0.95) water-immersion objective under spinning-disk confocal mode. Photodegradation was performed using a 405 nm solid-state laser (~7 mW at the sample plane) with a 40× (NA 1.15) water-immersion objective under spinning-disk confocal mode. Both imaging and photodegradation were controlled using NIS-Elements AR software (v5.30.04, Nikon).

Photodegradable mouse brain tissue samples were expanded in pure water and immobilized onto a 6-well glass-bottom plate (Cellvis) using the poly-L-lysine mounting method.[4] The expanded samples were first imaged using the 488 nm and 640 nm channels with a Z-step of 0.6 μm and the 20× objective on the above-mentioned confocal microscope. For photodegradation, the central field of view (FOV) was illuminated using the 405 nm laser and the 40× objective for several minutes until degradation was observed. After illumination, the same FOV was re-imaged using the 488 nm and 640 nm channels with a Z-step of 0.6 μm and the 20× objective.

#### 5.3 Expansion isotropy analysis

Regions of interest from the pre- and post-expansion images were selected and registered using a custom MATLAB code, as previously described.[5] The root mean square (r.m.s.) error was then calculated from all point-to-point distance measurements to evaluate expansion isotropy.

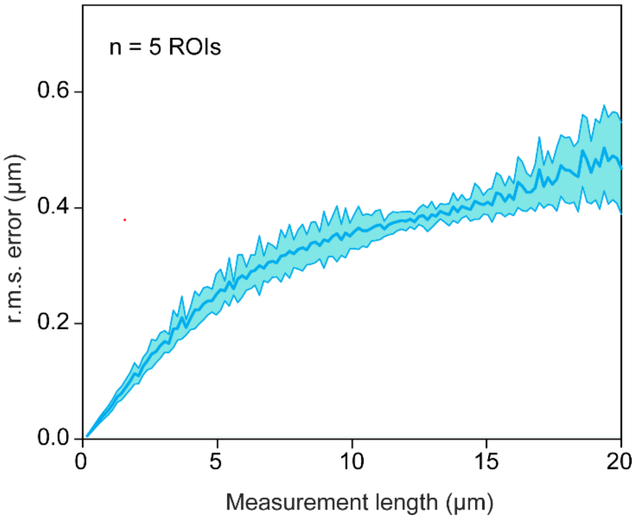


**Figure S1.** Expansion isotropy analysis of the PC-1500 gel. Root-mean-square (r.m.s.) expansion errors were measured for ~50 µm-thick mouse brain slices embedded and expanded with the PC-1500 gel. The blue line represents the mean, and the shaded cyan area represents the standard deviation across five regions of interest (ROIs; *n* = 5).

### 6. Rheometry

#### 6.1 Rheological measurements

Oscillatory rheological measurements were performed using an MCR 302 rheometer (Anton Paar) equipped with a 25 mm steel parallel-plate geometry. All measurements were performed at 25 °C using fully swollen PC-gels. Three independently prepared gels were measured for each formulation.

Before sample loading, a Petri dish was positioned around the lower measuring plate and secured to the rheometer stage with laboratory tapes to prevent movement during testing. The swollen gel sample was placed centrally between the parallel plates, and the upper plate was lowered until a normal force of 0.05 N was reached. Excess gel extending beyond the plate edge was removed. A fixed volume of 4 mL deionized water was then added to the surrounding Petri-dish reservoir to maintain sample hydration and minimize changes in the measured moduli caused by water loss or displacement during the measurement.

Frequency sweeps were performed first over an angular-frequency range of 0.1-10 rad/s at a fixed shear strain of 0.01%, using 20 logarithmically spaced measurement points. The low strain was selected to minimize strain-dependent softening and sample slippage during the frequency sweep. Amplitude sweeps were subsequently performed over a shear-strain range of 0.01-50% at a fixed angular frequency of 10 rad/s, using 20 logarithmically spaced measurement points. These measurements were used to assess the low-strain viscoelastic response and the onset of strain-dependent softening.

The storage modulus ($G'$) and loss modulus ($G''$) were recorded by the Anton Paar RheoCompass software and exported as CSV files for plotting and analysis.

#### 6.2 Reproducibility of oscillatory rheological measurements

To evaluate the reproducibility of the rheological measurements, three PC-gel replicates were independently prepared in separate gelation experiments for each formulation. Each gel was measured separately under the same conditions described in Section **6.1**. In short, these datasets represent independent gel preparations rather than repeated measurements of the same sample.

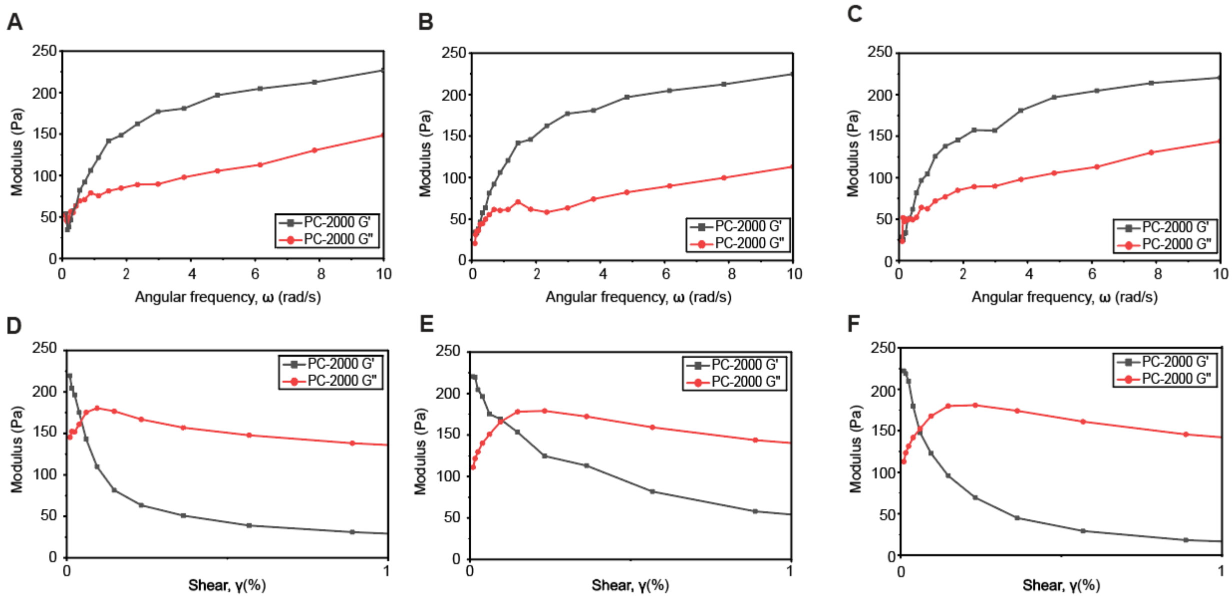


**Figure S2.** Oscillatory rheological characterization of swollen PC-2000 gels. **(A–C)** Frequency-sweep results and **(D–F)** amplitude-sweep results from three independently prepared PC-2000 gels. The storage modulus (G′) and loss modulus (G″) are shown for each sample. Measurement conditions are described in Section **6.1**. For clarity, the amplitude-sweep plots display the low-strain region from 0 to 1% shear strain.

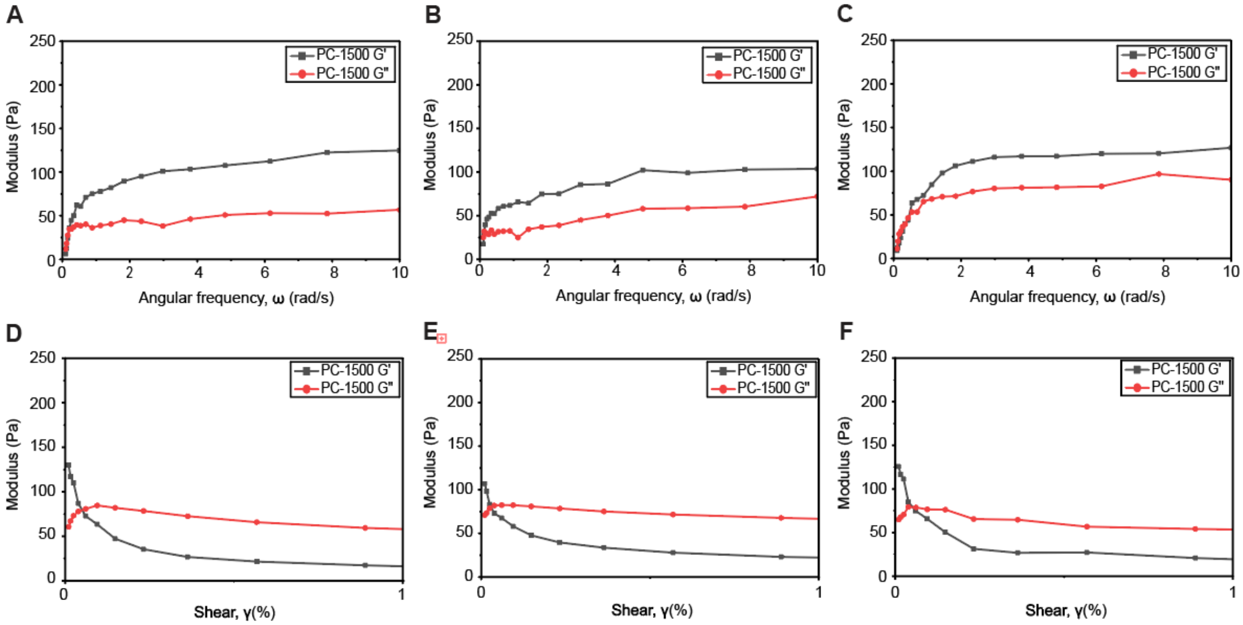


**Figure S3.** Oscillatory rheological characterization of swollen PC-1500 gels. **(A‑C)** Frequency-sweep results and **(D‑F)** amplitude-sweep results from three independently prepared PC-1500 gels. The storage modulus (G′) and loss modulus (G″) are shown for each sample. Measurement conditions are described in Section **6.1**. For clarity, the amplitude-sweep plots display the low-strain region from 0 to 1% shear strain.

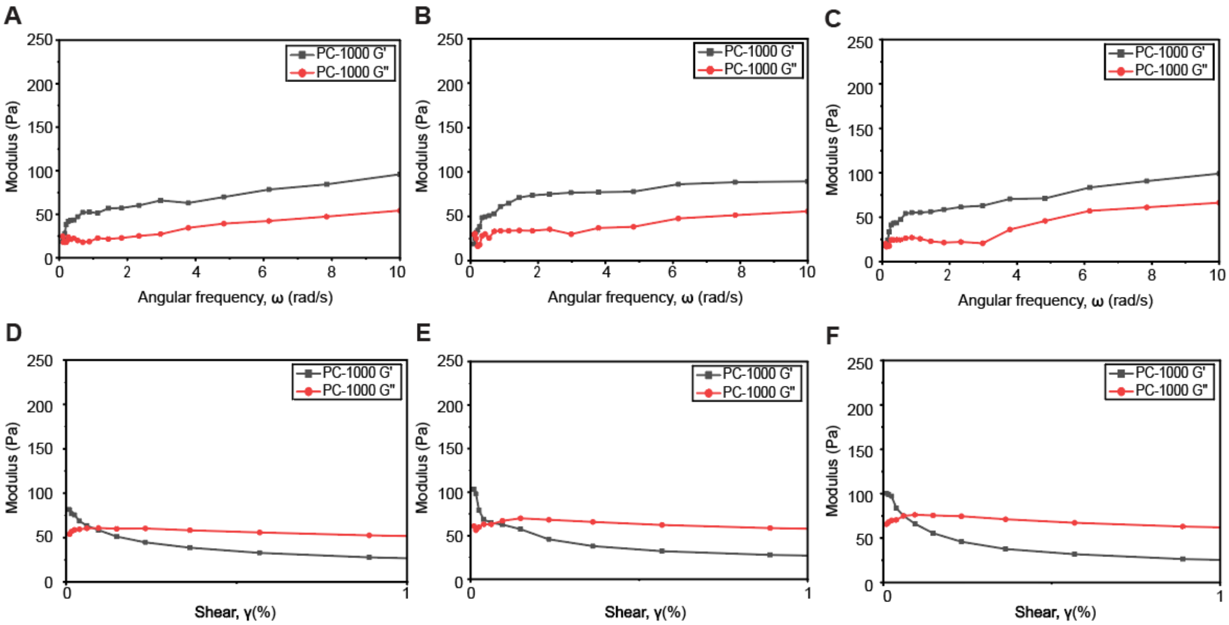


**Figure S4.** Oscillatory rheological characterization of swollen PC-1000 gels. **(A‑C)** Frequency-sweep results and **(D‑F)** amplitude-sweep results from three independently prepared PC-1000 gels. The storage modulus (G′) and loss modulus (G″) are shown for each sample. Measurement conditions are described in Section **6.1**. For clarity, the amplitude-sweep plots display the low-strain region from 0 to 1% shear strain.

### 7. Quantitative NMR analysis

Quantitative $^1H$ NMR (qNMR) was used to normalize the effective concentration of as-synthesized PC derivatives prior to gelation and to support fair comparison across PEG backbone lengths.[6,7] Because PEG-containing products can exhibit batch-to-batch purity variation and residual PEG-derived impurities, isolated mass alone was not used as the sole basis for cross-linker molarity.

#### 7.1 $^1H$ qNMR analysis

$^1H$ qNMR was performed to normalize the effective concentration of as-synthesized PC derivatives with varied PEG backbone lengths (**PC-2000, PC-1500, and PC-1000**) for the subsequent gelation experiments. In each experiment, cyclohexane (1 µL) was added to $CDCl_3$ (0.7 mL) as an internal standard, followed by dissolution of the weighed as-synthesized PC derivatives (**PC-2000**: 149.8 mg; **PC-1500**: 132.9 mg; **PC-1000**: 83.5 mg). Additional $D_2O$-based qNMR measurements were performed using acetone (1 µL in 0.7 mL $D_2O$) as the internal standard. The qNMR results obtained from the two solvent systems were consistent and used to assign the effective PC stock solution molarities. $^1H$ NMR spectra were acquired on a 500 MHz spectrometer (Bruker) and processed using standard phasing and baseline correction prior to integration. The cyclohexane resonance was integrated and normalized to 12.0 to account for its 12 equivalent protons ($N_{IS}$ = 12). A diagnostic acrylate vinyl resonance from the diacrylate-terminated PC derivatives was integrated as the analyte signal and converted to molar amount using the corresponding proton stoichiometry. Here, $N_A$= 2 per molecule, because the selected acrylate resonance represents one vinyl proton per acrylate terminus and each precursor bears two acrylate termini. The molar concentration of as-synthesized PC derivatives in each NMR tube was obtained from the integral ratio according to

$$n_{PC} = n_{IS} \times \frac{(I_A/N_A)}{(I_{IS}/N_{IS})}$$

where $n_{IS}$ is the known molar concentration of the internal standard. The resulting qNMR-derived molar concentrations of the as-synthesized PC derivatives were then used to scale the mass concentrations of aqueous PC derivative stock solutions so that an identical molar amount of PC derivatives was used for gelation. **PC-2000** stock was defined as the reference (6.25 mg/100 µL), and the target weight concentrations for **PC-1500** and **PC-1000** were adjusted using qNMR results to match the reference molar concentrations. These normalized stock solutions were then used for PC-gel formation under otherwise identical conditions, ensuring that observed differences in gel properties arise from differences in the cross-linker backbone length rather than unequal concentrations. The qNMR-derived molar concentrations of the as-synthesized PC derivatives were used to normalize the stock solutions for gel preparation. **PC-2000** was used as the reference, and the target stock concentrations for other PC derivatives were adjusted based on qNMR results to ensure the same stock volume delivered the same effective molar amount of cross-linker during polymerization.

### 8. NMR spectra

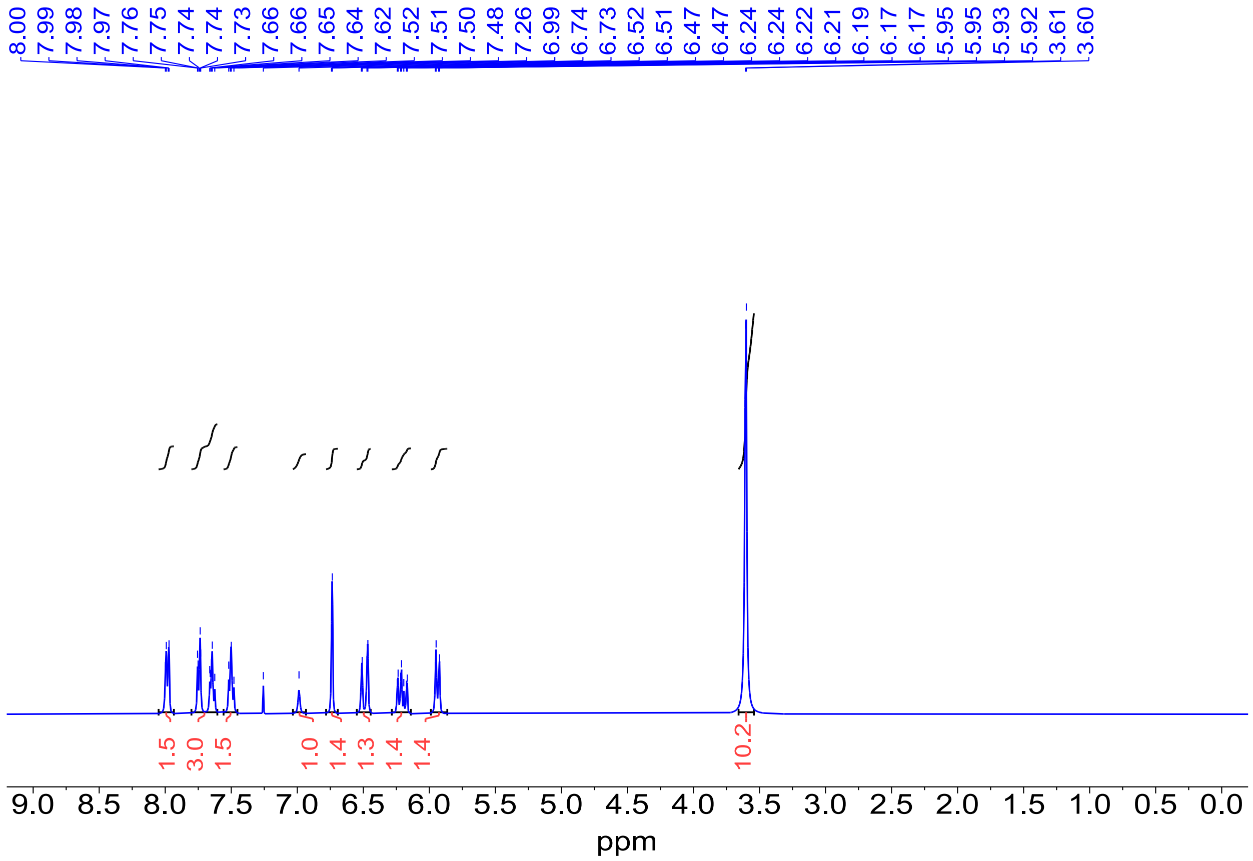


**Figure S5.** [1]*H* NMR spectrum (500 MHz) of **PC-200** in $CDCl_3$. *: Residual solvent peak.

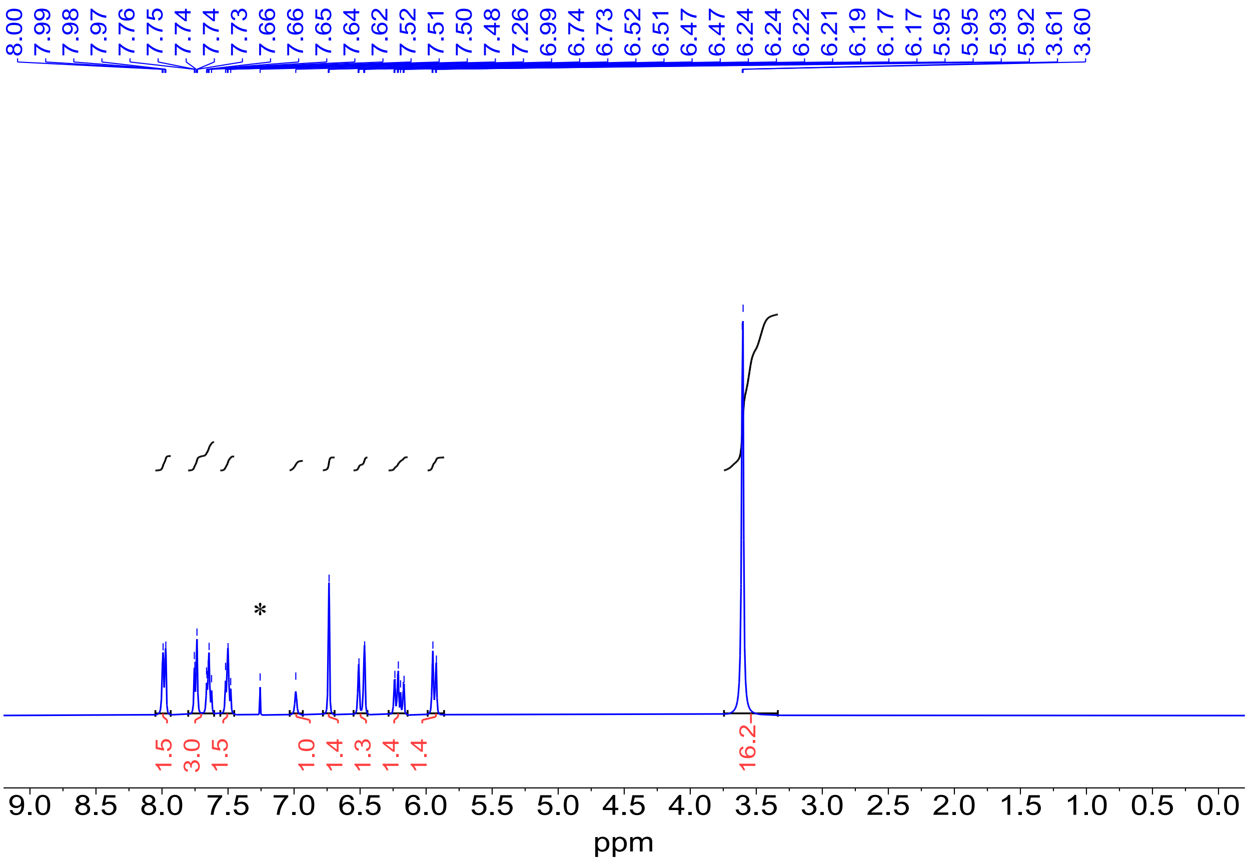


**Figure S6.** $^{1}H$ NMR spectrum (500 MHz) of **PC-500** in $CDCl_3$. *: Residual solvent peak.

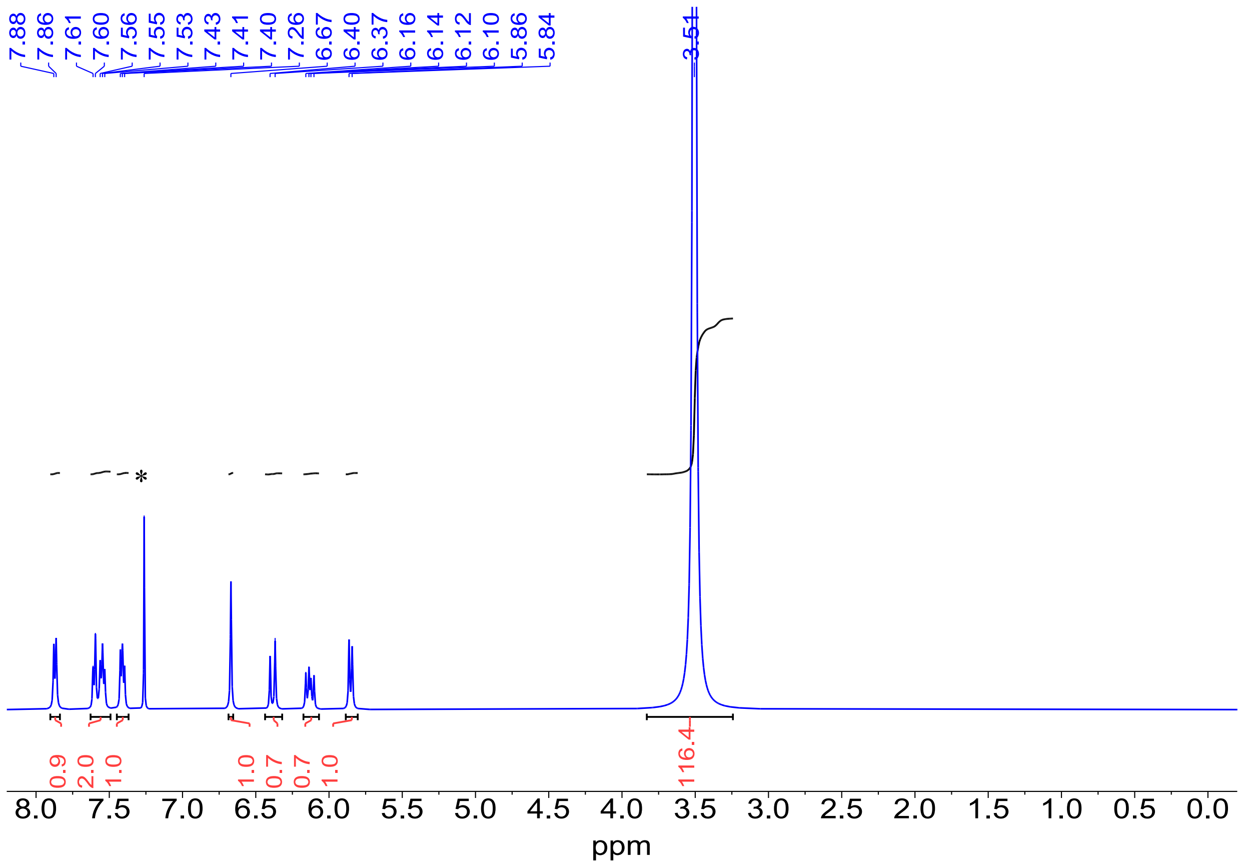


**Figure S7.** $^{1}H$ NMR spectrum (500 MHz) of **PC-1000** in $CDCl_3$. *: Residual solvent peak.

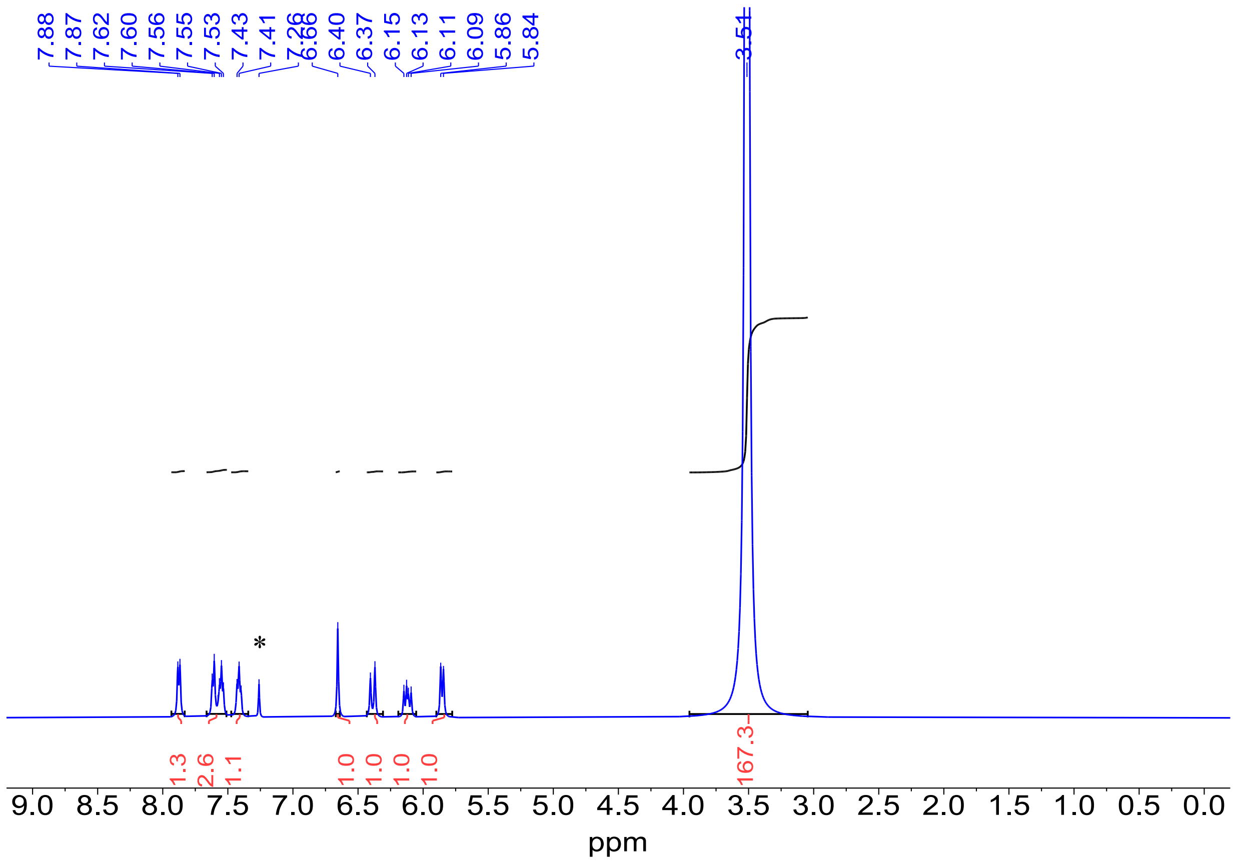


**Figure S8.** $^{1}H$ NMR spectrum (500 MHz) of **PC-1500** in $CDCl_3$. *: Residual solvent peak.

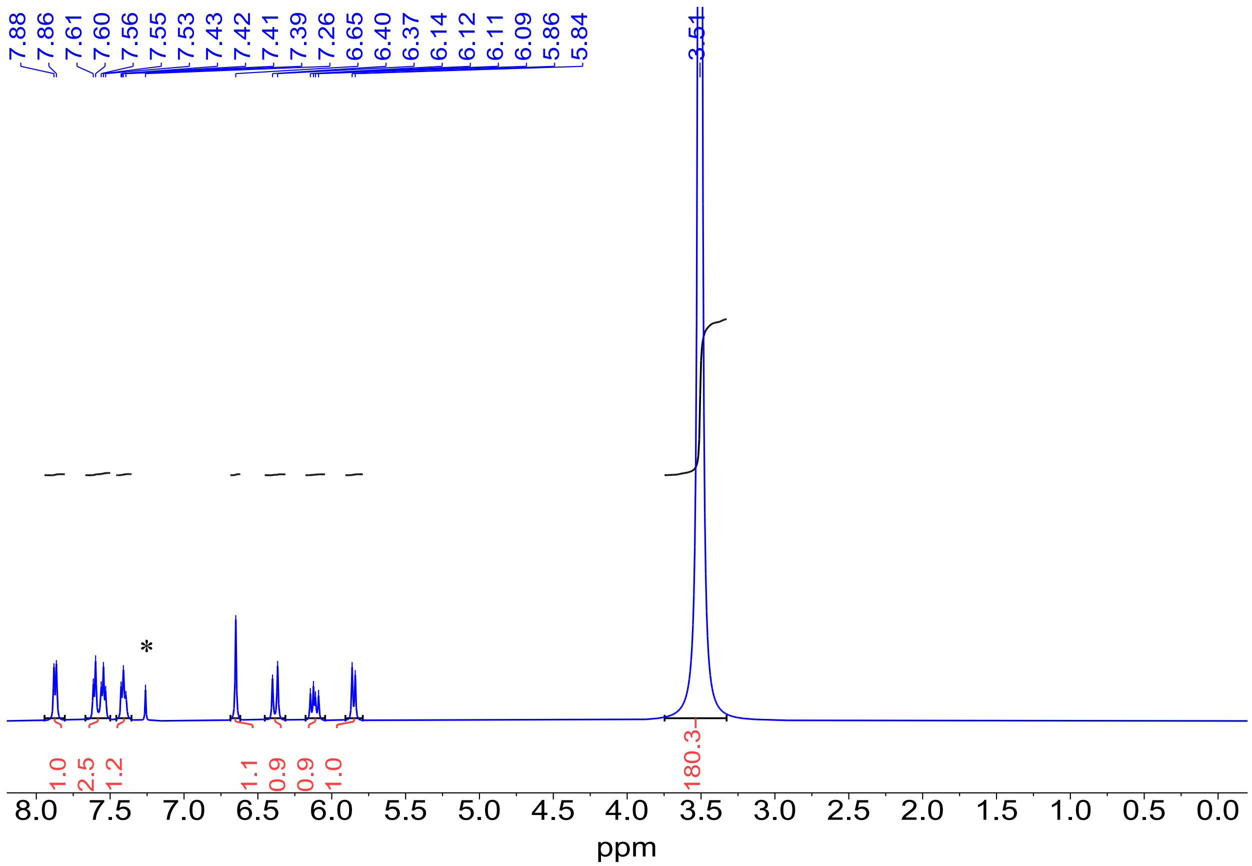


**Figure S9.** $^{1}H$ NMR spectrum (500 MHz) of **PC-2000** in $CDCl_3$. *: Residual solvent peak.

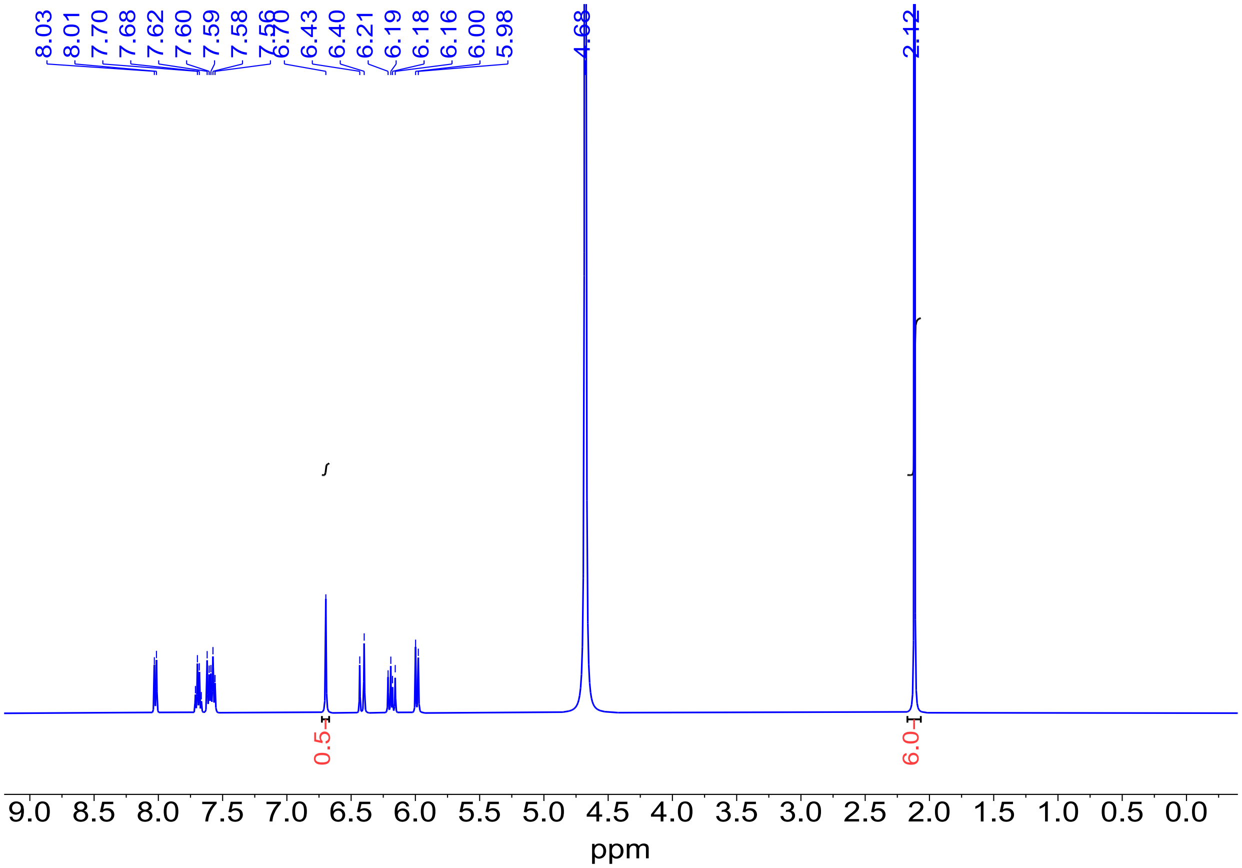


**Figure S10.** Quantitative ${}^{1}H$ NMR (qNMR) analysis of **PC-200** in $D_2O$.

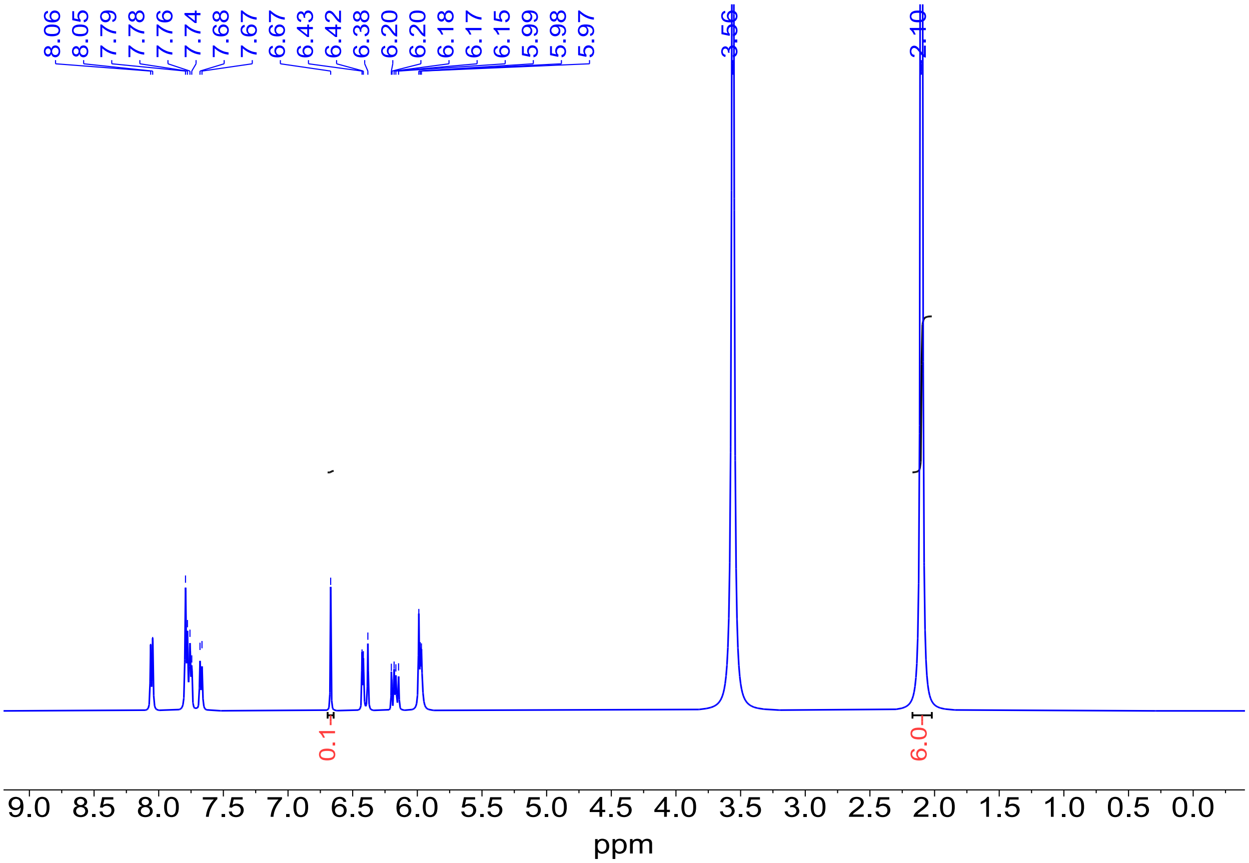


**Figure S11.** Quantitative $^{1}H$ NMR (qNMR) analysis of **PC-500** in $D_2O$.

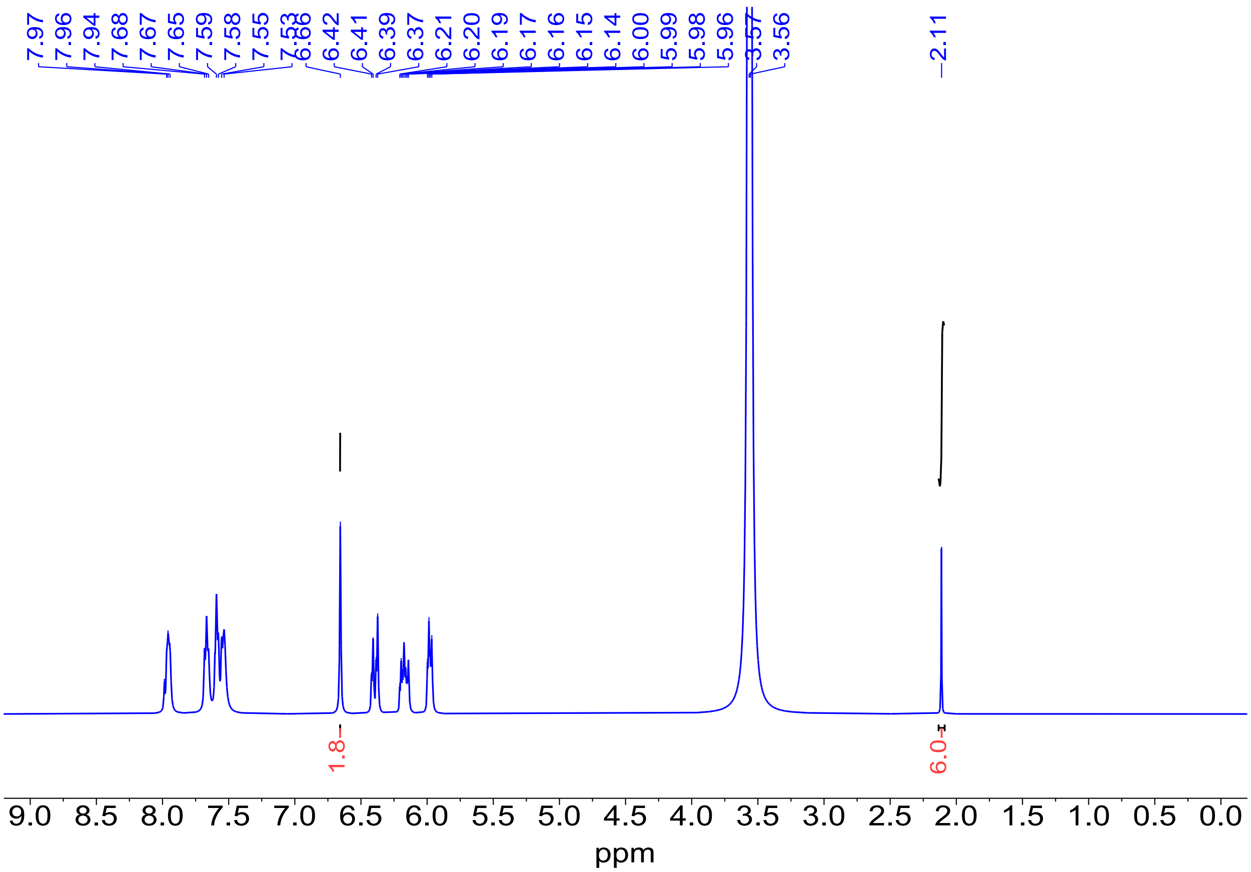


**Figure S12.** Quantitative $^{1}H$ NMR (qNMR) analysis of **PC-1000** in $D_2O$.

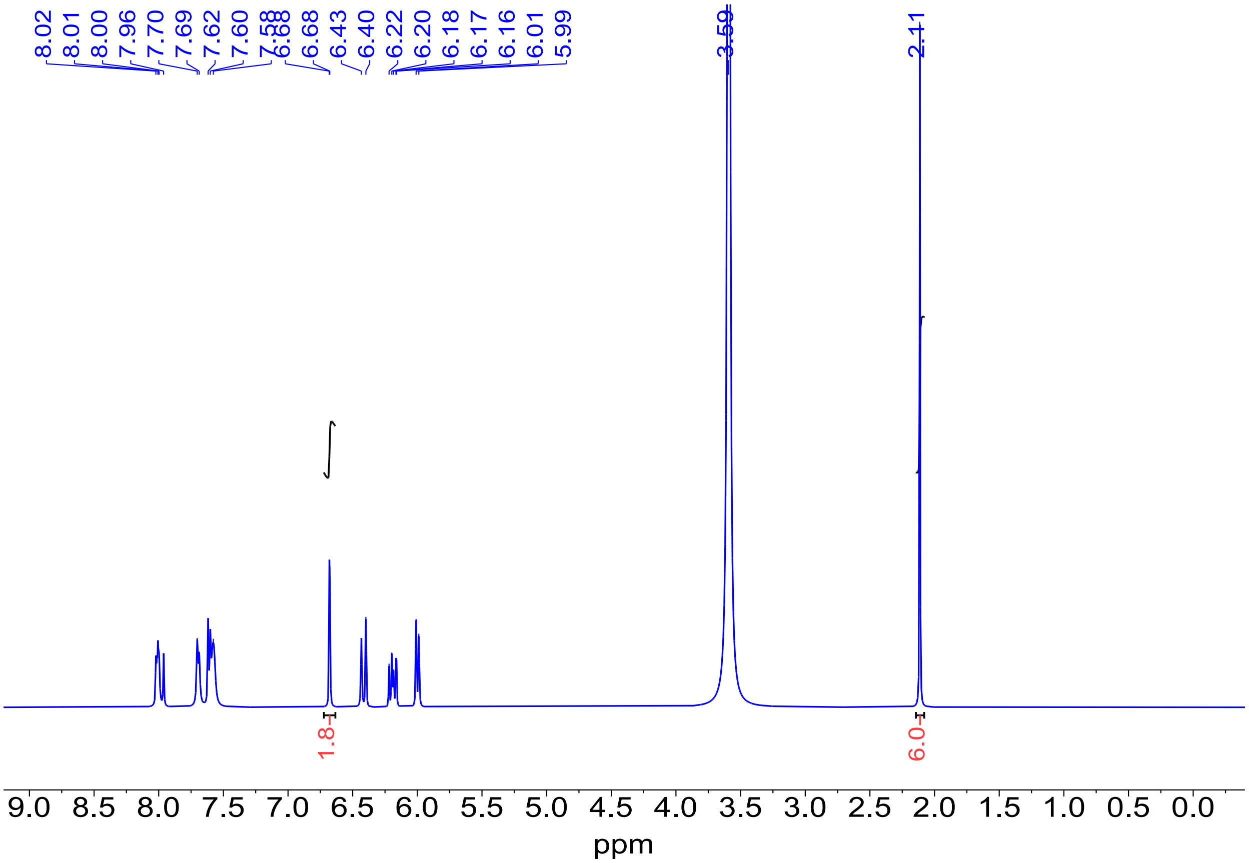


**Figure S13.** Quantitative $^{1}H$ NMR (qNMR) analysis of **PC-1500** in $D_2O$.

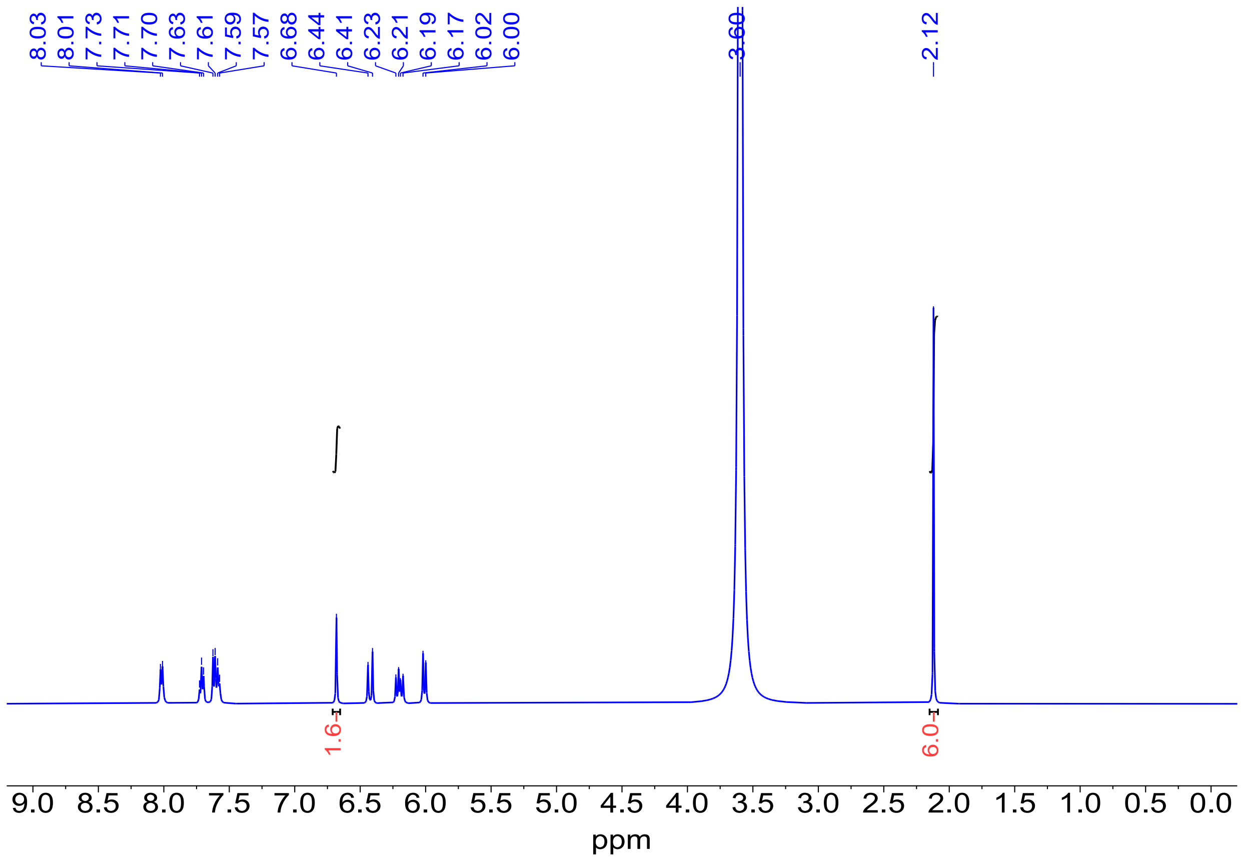


**Figure S14.** Quantitative $^{1}H$ NMR (qNMR) analysis of **PC-2000** in $D_2O$.

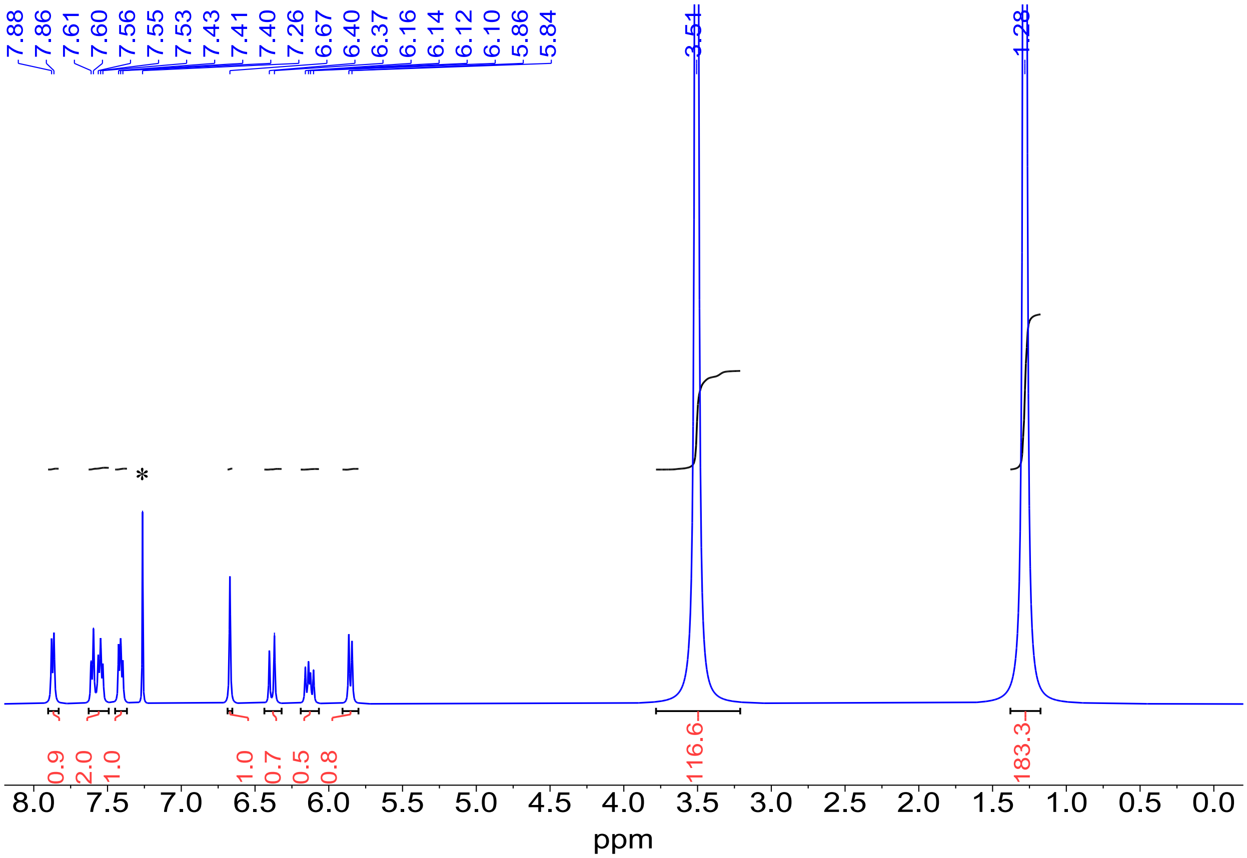


**Figure S15.** Quantitative $^{1}H$ NMR (qNMR) analysis of **PC-1000** in $CDCl_3$. *: Residual solvent peak.

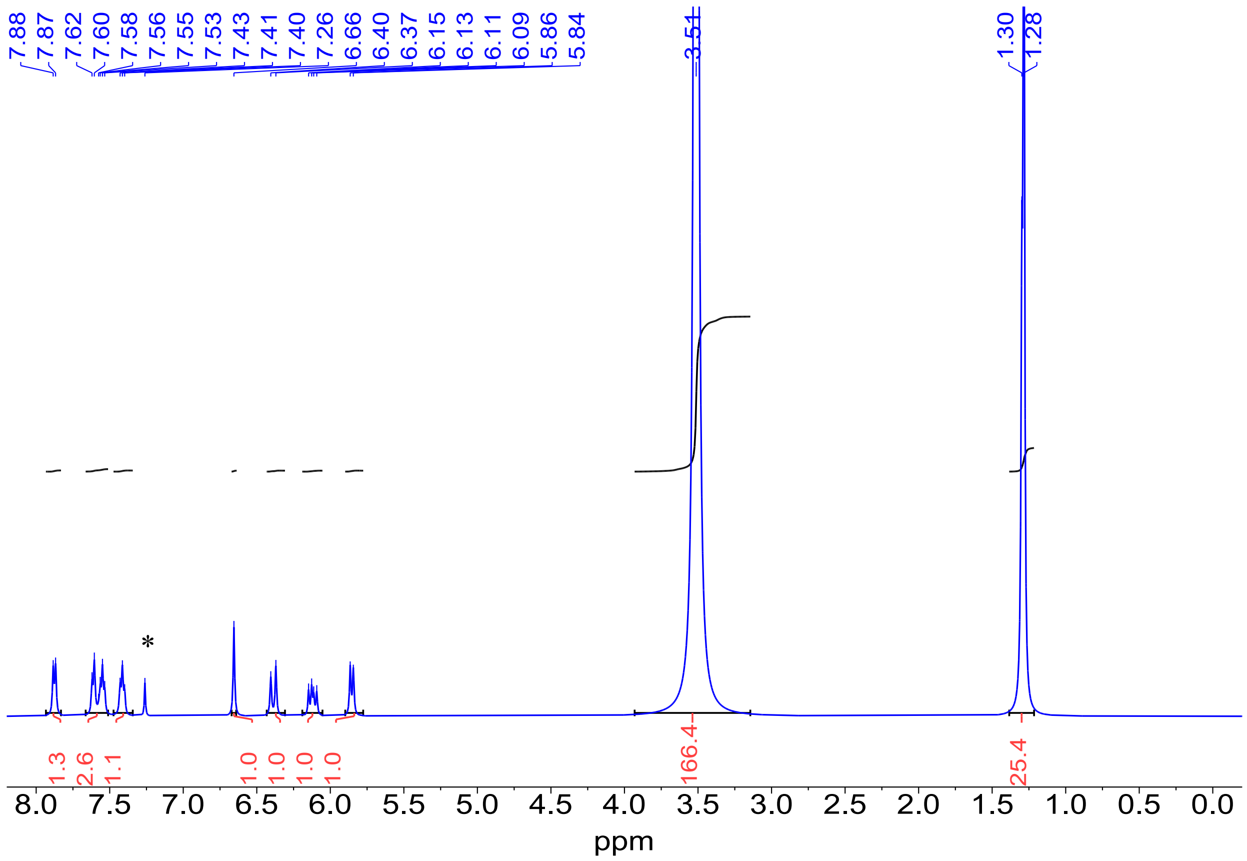


**Figure S16.** Quantitative [1]*H* NMR (qNMR) analysis of **PC-1500** in $CDCl_3$. *: Residual solvent peak.

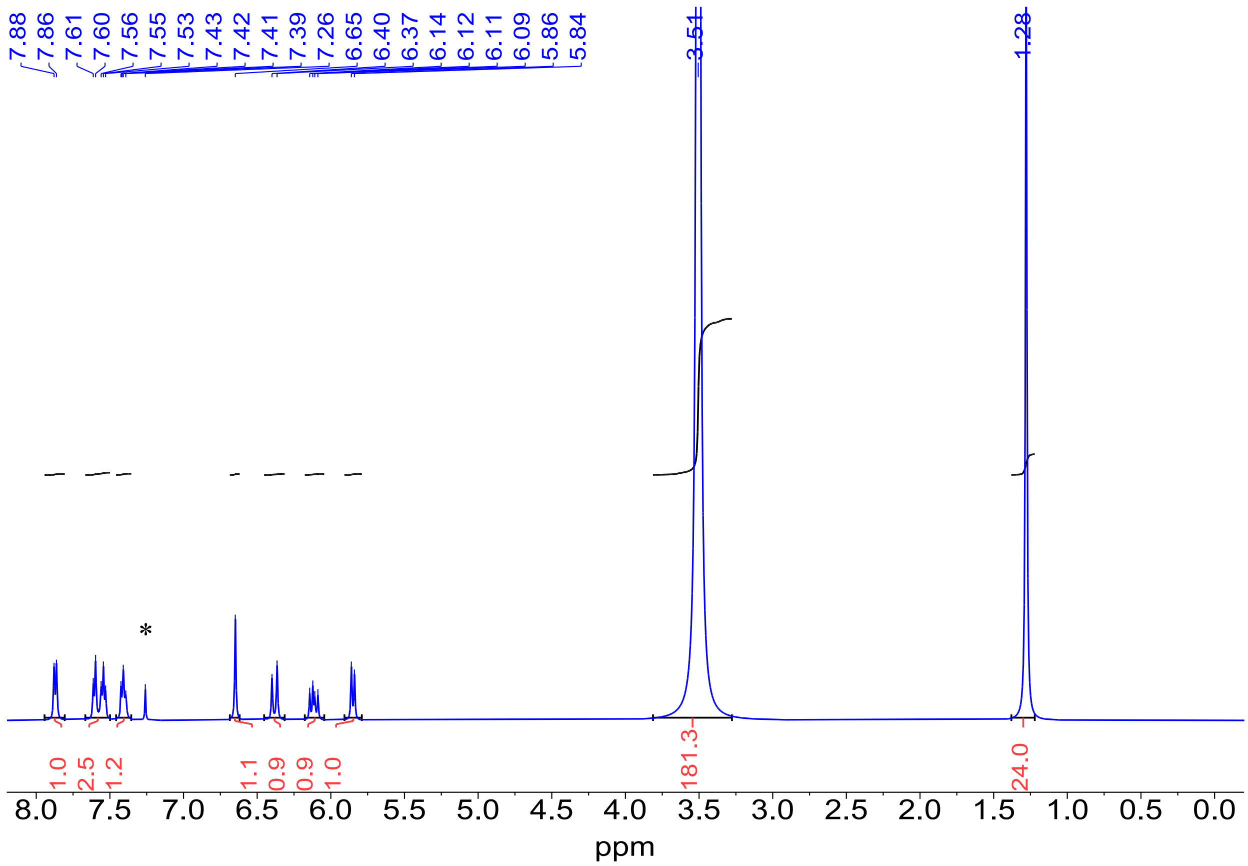


**Figure S17.** Quantitative $^{1}H$ NMR (qNMR) analysis of **PC-2000** in $CDCl_3$. *: Residual solvent peak.
